# Normalization by orientation-tuned surround in human V1-V3

**DOI:** 10.1101/2021.11.06.467486

**Authors:** Zeming Fang, Ilona Bloem, Catherine Olsson, Wei Ji Ma, Jonathan Winawer

**Author notes:** Corresponding author: Zeming Fang.

## Abstract

An influential account of neuronal responses in primary visual cortex is the normalized energy model. This model is often implemented as a two-stage computation. The first stage is the extraction of contrast energy, whereby a complex cell computes the squared and summed outputs of a pair of linear filters in quadrature phase. The second stage is normalization, in which a local population of complex cells mutually inhibit one another. Because the population includes cells tuned to a range of orientations and spatial frequencies, the result is that the responses are effectively normalized by the local stimulus contrast. Here, using evidence from human functional MRI, we show that the classical model fails to account for the relative responses to two classes of stimuli: straight, parallel, band-passed contours (*gratings*), and curved, band-passed contours (*snakes*). The snakes elicit fMRI responses that are about twice as large as the gratings, yet a traditional divisive normalization model predicts responses that are about the same. Here, motivated by these observations and others from the literature, we implement a divisive normalization model, in which there is a preferential inhibition between cells matched in orientation tuning (“tuned normalization”). We first show that this model accounts for differential responses to these two classes of stimuli. We then show that the model successfully generalizes to other band-pass textures, both in V1 and in extrastriate cortex (V2 and V3). We conclude that even in primary visual cortex, complex features of images such as the degree of heterogeneity, can have large effects on neural responses.

## 1. Introduction

Primary visual cortex (“V1”) has served as a testing ground for studying physiology, anatomy, brain development, neuroimaging, and computational modeling. There has been considerable success in developing general model forms that capture many of the encoding properties of V1 neurons reasonably well over a range of stimulus conditions, including the normalized energy model of V1 complex cells (<u>Heeger 1993</u>). This type of model, like many others (reviewed in chapter 6 of <u>Wandell</u> <u>1995</u>; <u>Hubel and Wiesel 1962</u>), includes a linear filter as the first stage, i.e., a weighted sum of the stimulus intensity over space and time. In a second stage, the outputs of the filter are squared and summed across nearby spatial locations or across phase (<u>Adelson and Bergen 1985</u>; <u>Pollen and Ronner</u> <u>1983</u>; <u>Heeger 1992a</u>). If the outputs are summed across a pair of linear filters tuned to the same frequency, orientation, and location, but differing in phase by 90 deg, it is called an energy model. In the third stage, the response of each neuron is normalized (divisively suppressed) by the (un-normalized) outputs of the nearby neural population (<u>Heeger 1992b</u>, <u>1993</u>). This effectively adjusts the gain based on the contrast energy in the image or image patch. There is substantial evidence that each of these three operations–linear filtering, energy, and normalization–contribute to the responses of V1 neurons (<u>reviewed by Carandini et al. 2005</u>).

The normalized contrast energy model, though initially developed to explain the outputs of single neurons, has also been successfully applied to functional MRI data in human visual cortex. First, a contrast energy model without normalization, applied to voxels in V1, V2, and V3, was used to predict BOLD responses (encoding) and to infer the viewed images from the BOLD responses (decoding) (<u>Kay et</u> <u>al. 2008</u>). Subsequent work showed that incorporating a normalization-like non-linearity improved model accuracy when testing stimuli that varied substantially in size (<u>Kay, Winawer, Mezer, et al. 2013</u>) or pattern (<u>Kay, Winawer, Rokem, et al. 2013</u>), and that normalization could account for the BOLD contrast response function for gratings with and without masking stimuli at other orientations (<u>Brouwer</u> <u>and Heeger 2011</u>). These models have also shown good prediction accuracy for similar stimulus sets used in human intracranial electrode recordings of visual cortex (<u>Hermes et al. 2019</u>; <u>Winawer et al.</u> <u>2013</u>).

The normalized contrast energy model, although successful at accounting for responses to a range of stimuli, nonetheless fails to explain some phenomena. There is some evidence that the standard model declines in accuracy for natural images (<u>David, Vinje, and Gallant 2004</u>; <u>Olshausen and Field 2005</u>; <u>Vinje</u> <u>and Gallant 2000</u>) (<u>but see alsoRust and Movshon 2005</u>). Even testing with simple patterns, early studies of V1 and extrastriate electrophysiology showed that some cells had tuning properties differing from simple or complex cells, called “hypercomplex” cells, many of which were associated with “end- stopping” {Hubel, 1968 #3338}{Hubel 1965}. Recent V1 two-photon calcium recordings included a large stimulus set and found that many cells, with or without end-stopping, were surprisingly sparsely tuned, often sensitive to complex patterns such as crosses or composite features (<u>Tang et al. 2018</u>). It is unlikely that the standard energy model would predict the kind of tuning they observed, although a model was not fit to the data. Other studies with single unit electrophysiology found that a normalization model could be successful but only if the normalization was flexible, such that its strength depended on statistical dependencies in the image (<u>Coen-Cagli, Kohn, and Schwartz 2015</u>). In human fMRI studies, the responses to natural/complex images in V1 also appear to be influenced by statistical dependencies and image context (<u>Mannion, Kersten, and Olman 2015</u>; <u>Qiu et al. 2016</u>), factors unlikely to influence the predictions of the normalized energy model. Even in relatively simple artificial images with a fixed amount of total contrast energy, the BOLD response is lower when there is a single orientation compared to when there are two divergent orientations (<u>Bloem and Ling 2019</u>; <u>Klimova, Bloem, and Ling</u> <u>2021</u>).

In visual areas beyond V1, the normalized energy model is expected to be incomplete, as circuits in these areas contribute new computations. There are no widely adopted standard encoding models for these areas analogous to the normalized energy model for V1, but there has been some success in modeling patterns in the extrastriate responses by incorporating higher-order statistical dependencies of the modeled V1 outputs (<u>Freeman et al. 2013</u>; <u>Movshon and Simoncelli 2015</u>; <u>Okazawa, Tajima, and</u> <u>Komatsu 2017</u>) or sensitivity to second-order contrast (<u>Kay, Winawer, Rokem, et al. 2013</u>).

Here, using evidence from human functional MRI, we show that the classical model fails to account for the relative responses to two classes of stimuli: straight, parallel, band-passed contours (*gratings*), and curved, band-passed contours (*snakes*) (**Figure 1**). The snakes elicit fMRI responses that are about twice as large as the gratings, yet traditional energy models, including normalized energy models, predict responses that are about the same. This is a large model failure which, in conjunction with the other failures of the simple normalization model described above, motivated us to implement a model in which the normalization is tuned, meaning the normalization pool for a given neural channel has the same orientation tuning as the channel being normalized. We also developed and implemented a computational model that achieves tuned normalization in a different way, in which responses are normalized not by the sum of the contrast energy, but by the anisotropy (standard deviation) in contrast energy computed across orientation channels. Both models account for the differences in responses to snakes vs gratings, supporting the proposal that normalization depends on the spatial arrangement of image features, and not just the total amount of contrast energy.

**Figure 1.**
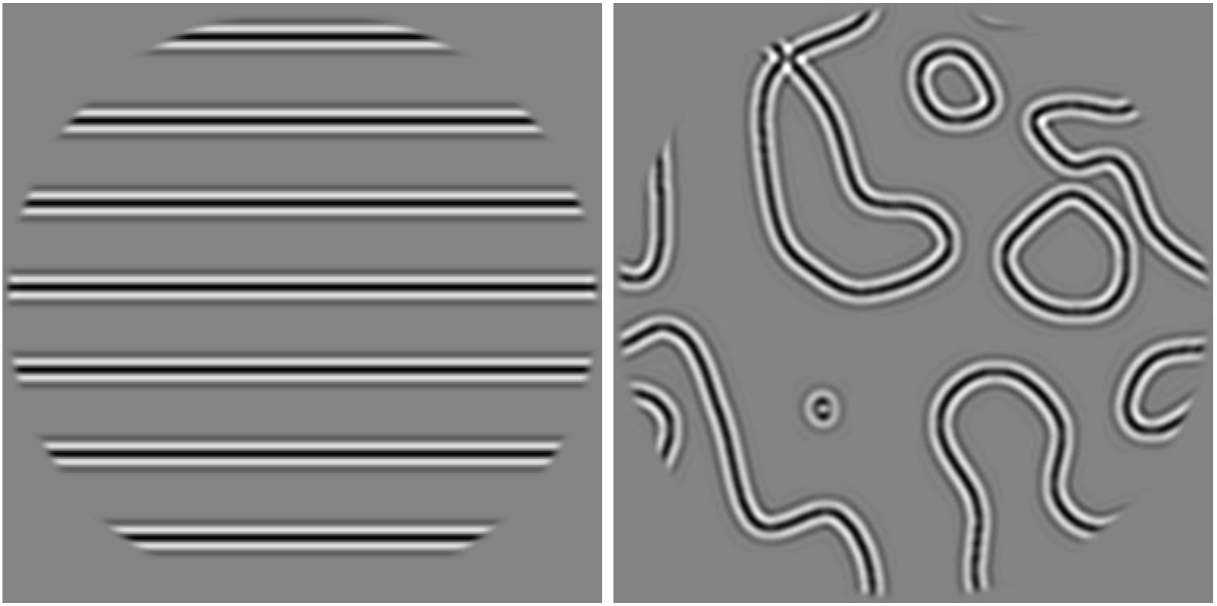
Example straight-line and curved-line stimuli from our experiments. We observe that in human V1, V2, and V3, stimuli with long straight lines (gratings) reliably evoke a smaller fMRI response than similar stimuli with curved lines.

## 2. Results

### 2.1. The fMRI BOLD responses to curved patterns are larger than to straight patterns

We first consider an observation about fMRI responses to two classes of simple, grayscale, band-passed, static images. For one class, the stimuli contain several curved contours, which we refer to throughout as snakes. For the other class, the stimuli contain several straight, parallel contours, which we refer to as gratings. The surprising observation is that for V1, V2 and V3, the fMRI responses are substantially larger for the snakes than for gratings (**Figure 2**). The responses to the gratings, irrespective of density, are only about as high as the response to the sparsest snakes. We confirm this pattern with three additional fMRI data sets, which also show larger responses to snakes than gratings in V1, V2 and V3 (**Figure S2**).

**Figure 2.**
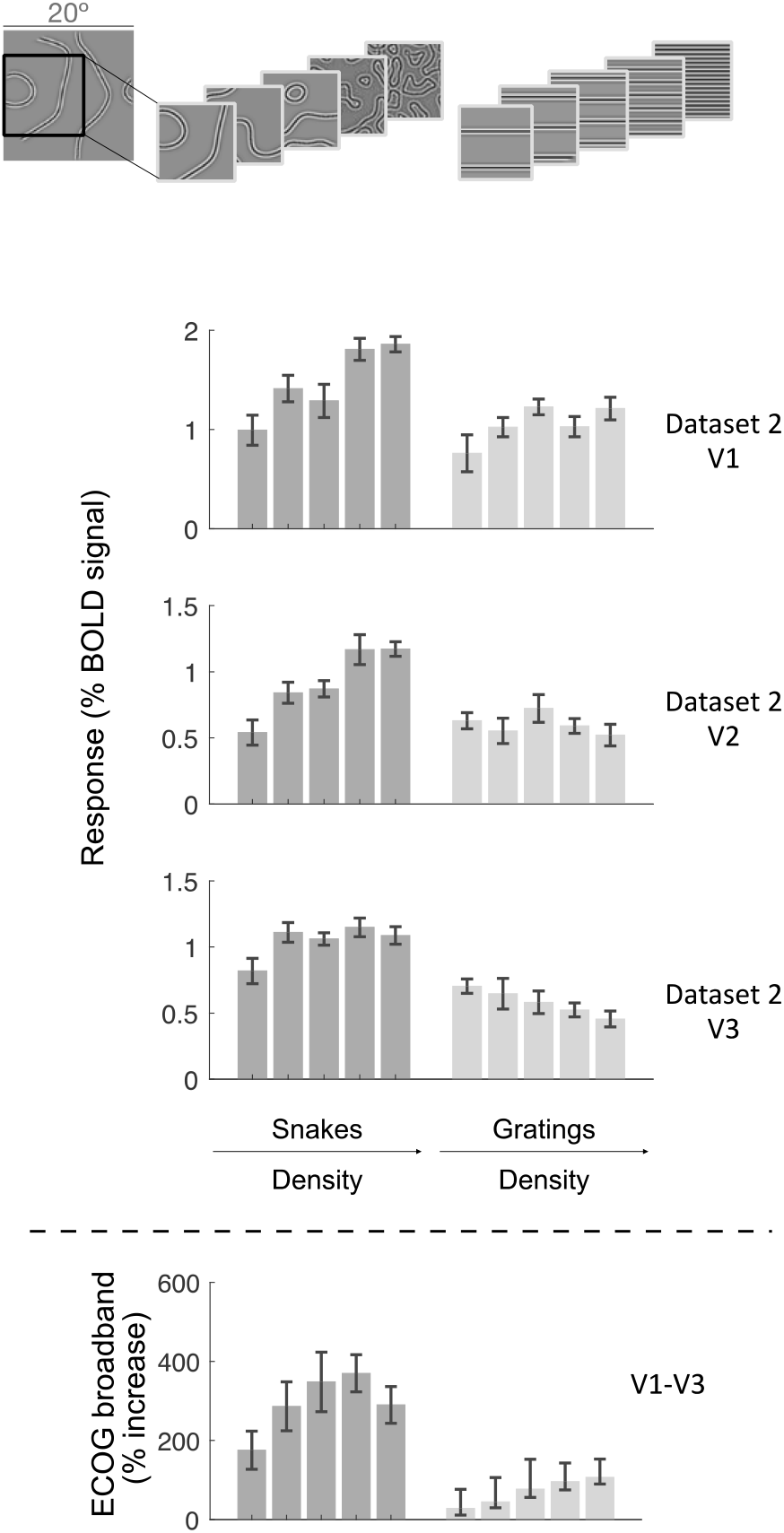
FMRI responses are larger for curved patterns than for straight patterns. The mean fMRI responses within a visual area are plotted for V1, V2, and V3 in dataset S2. For both curved stimuli (*snakes*, dark bars) and straight stimuli (*gratings*, light bars), data are plotted in order of increasing texture density from left to right. Examples of the 5 densities are shown above. Error bars are the standard deviation of the mean, bootstrapped across fMRI runs. Responses are larger for the *snakes* than *gratings*. The same effect is also observed for datasets S1, S3 and S4 (Supplementary Figure S2). Bottom panel: For comparison, we replot data from intracranial recordings (ECoG) from Hermes et al 2019, which also show larger responses for *snakes* than for *gratings*. The full set of stimuli is shown in Supplementary Figure S1. Data plotted from the function *s4_visualize(’figure2’)* in the <u>GitHub code</u> <u>repository</u>.

To check whether this pattern of results is specific to fMRI, we replot published intracranial data from Hermes et al. (<u>2019</u>), in which human subjects with ECoG recordings viewed a similar set of stimuli (**Figure 2**, bottom panel). The ECoG data, plotted as the percent increase in broadband power over baseline, show the same general effect as the fMRI data: The responses to snakes are much larger than to gratings, irrespective of texture density, indicating that the effect is not limited to the fMRI BOLD response. (See Hermes et al. (<u>2019</u>) for methodological details.)

In the next four sections, we describe the four models that are fit to the data. For simplicity, for each model we fit parameters to the aggregate (average) data within a visual area, rather than to each voxel individually. All the stimuli are texture-like, meaning that they have similar properties across the whole image aperture, and for each stimulus class, nine different exemplars were shown per 3-s trial. The different exemplars have the same higher-order statistics but vary in the precise spatial distributions. Hence model computations, such as contrast energy, would result in similar values for spatially localized portions of the image (as one would compute for an individual voxel) and for the whole image (as we compute to model the aggregate response of a visual area). For this reason, we did not include model parameters for the spatial location or spatial extent of the receptive fields for individual voxels. We first describe models fit to the two classes of stimuli described above (gratings and snakes) that vary in either density or contrast, which we refer to as *target stimuli*. We then summarize fits to the larger set of stimuli viewed by the subjects.

All four models consist of three components, *filtering*, *spatial pooling*, and *power-law nonlinearity*. In the filtering stage, we computed the contrast energy of the stimuli at 8 orientations and 4 spatial frequencies (see Methods for details). Although stimuli were designed to have power concentrated close to 3 cycles per degree, there is some spillover to lower and higher frequencies, which is why we use multiple spatial frequency bands in our models. The spatial pooling stage pools the contrast energy to yield a total contrast energy. In some models, the pooling stage also includes divisive normalization. The output of the pooling is a scalar, which is then passed through a power-law nonlinearity to predict BOLD amplitude in units of percent signal change. The power-law nonlinearity achieves compressive spatial summation (<u>Kay, Winawer, Mezer, et al. 2013</u>). All models have the same filtering (first stage) and output -linearity (third stage). They differ in the spatial pooling stage.

### 2.2. The larger response to snakes is not captured by a simple contrast energy model

A *standard contrast energy* model pools the contrast energy by simply summing it across space, orientations, and spatial frequencies to give a total contrast energy. It predicts that responses should increase with both stimulus contrast and with density of the pattern. This model does not predict a larger response to the snakes than to gratings, contrary to the data (**Figure 3**). In fact, the cross-validated variance explained is low (V1, V2) or even negative (V3) in the example data, meaning that the model prediction is less accurate than it would have been if it simply predicted the mean response across all stimuli. In short, the contrast energy model provides a poor fit to the fMRI data in V1-V3 for these classes of stimuli. It is also a poor fit to the target stimuli in the other three data sets (**Figure 8**, **Figure S2a**). This does not mean contrast energy models are always poor fits to fMRI responses in V1-V3. For example, when stimuli vary in how contrast energy is distributed across space, a contrast energy model can capture a lot of the variance in the responses across images, as shown by Kay et al. (<u>2008</u>).

**Figure 3:**
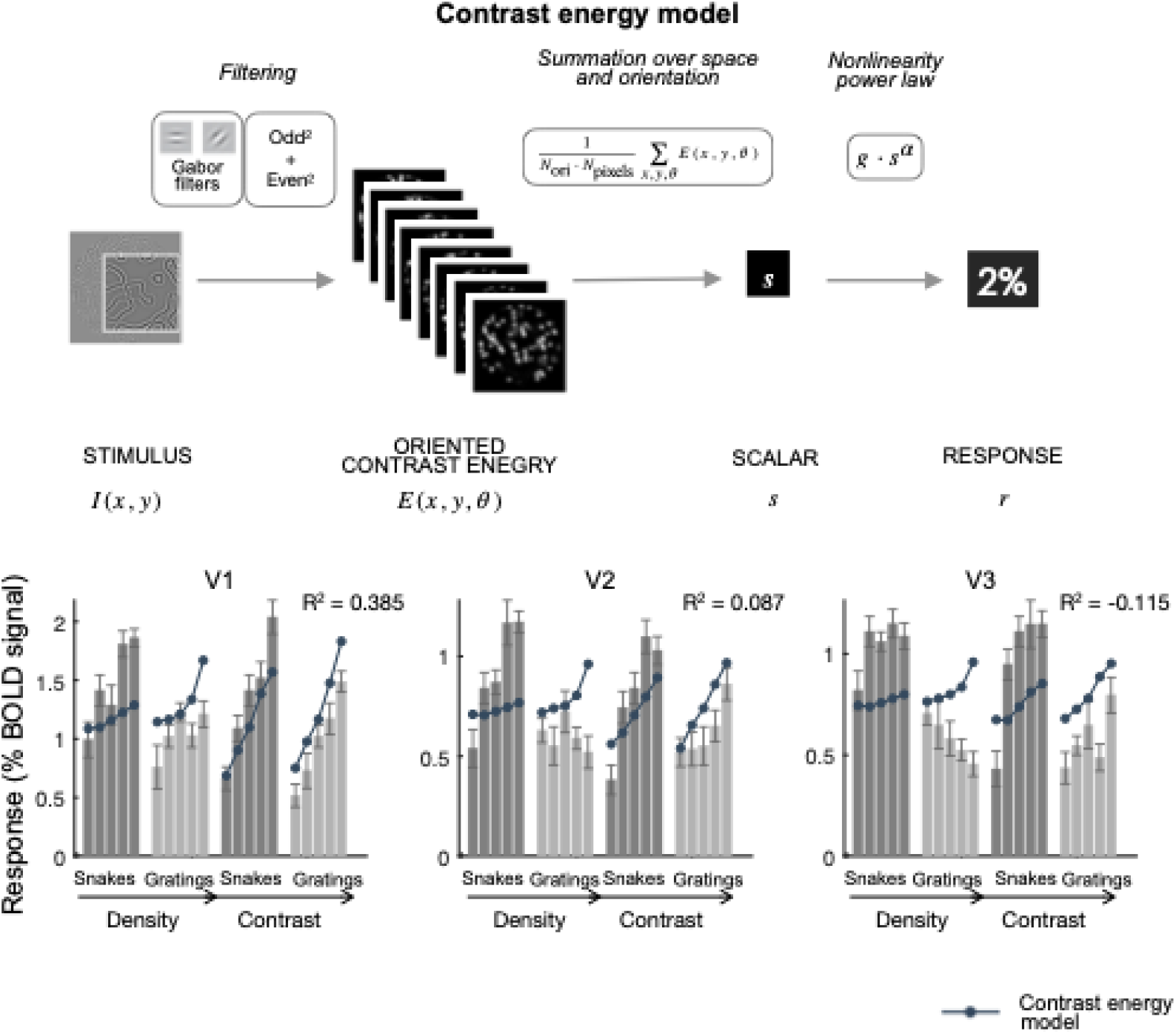
Contrast energy model does not account for V1-V3 responses to these stimuli. The mean fMRI responses from V1, V2, and V3 are shown for snakes and grating stimuli that vary in density and contrast for the subject (Dataset 2). The bar plots show the mean and standard error of the responses. The dots are the cross-validated predictions from the contrast energy model. Dark bars represent snake stimuli and light bars represent grating stimuli. Each group of stimuli is arranged in increasing order of either density or contrast. Data and model fits plotted from the function *s4_visualize(’figure3’)* in the <u>code repository</u>. The upper panel is a schematic representation of the contrast energy model. To simplify the schematic, we show E after pooling across spatial frequency bands. See *Figure S2a* for fits to all 4 datasets. See Table S5 for model parameters.

### 2.3. The larger response to snake stimuli is not captured by an untuned normalization model

We then added divisive normalization to the model. After computing contrast energy, we normalize the outputs by dividing the contrast energy at each pixel by the contrast energy of a normalization pool (**Figure 4, upper panel**). The normalization pool includes nearby locations, all spatial frequencies, and all orientations, giving it the name *untuned normalization model* (**Figure 4, lower left panel**).

**Figure 4.**
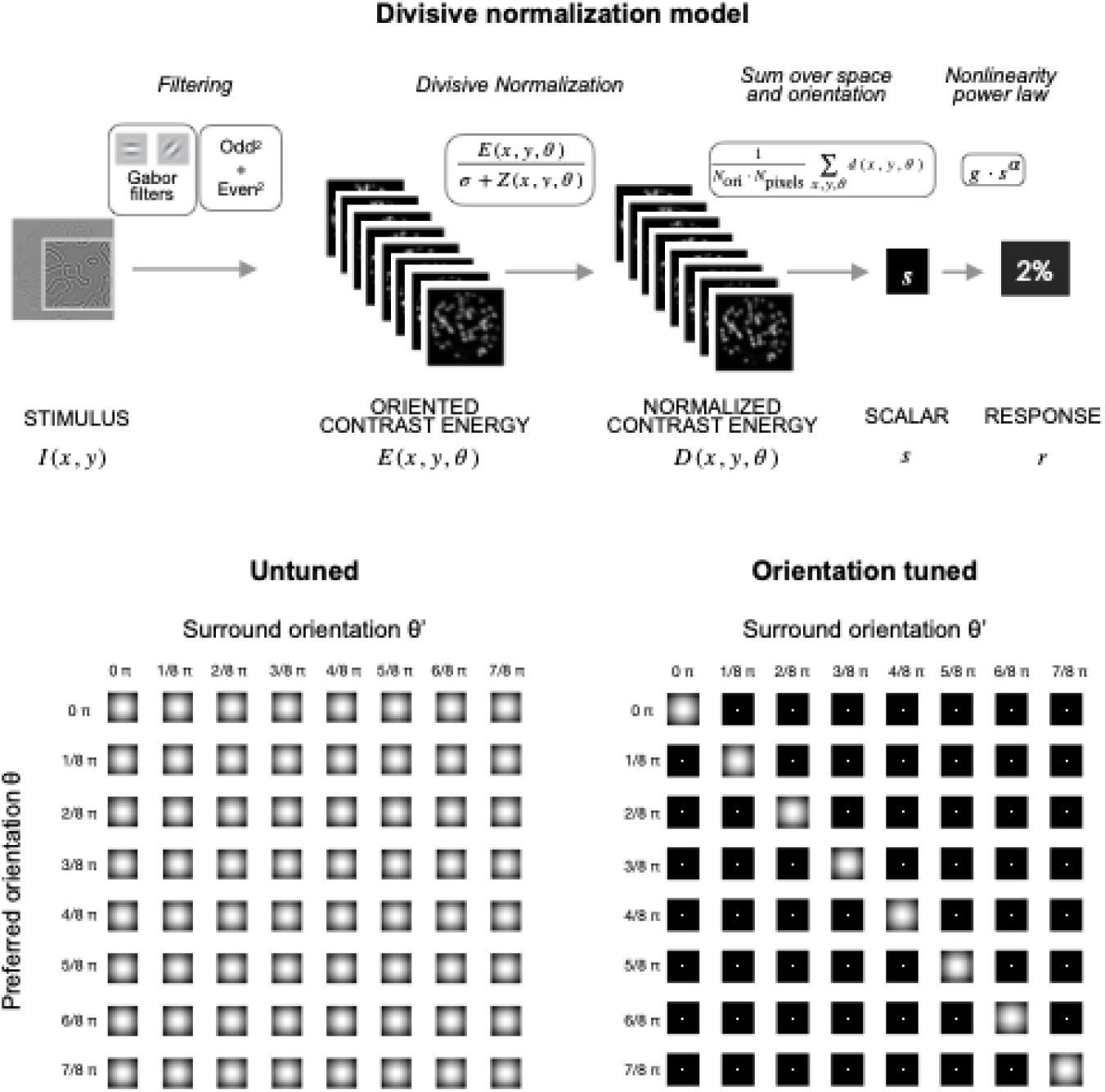
Two divisive normalization models. The upper panel indicates the process of the models. The lower panels are the weight filters used to calculate the normalization for divisive normalization variants. Filters are plotted from the function *s4_visualize(’figure4’)* in the <u>code repository</u>. See *Figure S2b-c* for fits to all 4 datasets.

The untuned normalization model and the contrast energy model have a similar pattern of results with similar variance explained. One difference between their predictions is that the normalization model results in more saturation at high contrast, as expected from divisive normalization (<u>Heeger 1992b</u>). This is especially evident in V2 and V3 at the highest stimulus contrast. The reason that the normalization model and the contrast energy model have a similar overall pattern of predictions is that the power law output nonlinearity, included all models, can do some of the work of normalization (Kay et al., 2013b, section “Relationship to Divisive Normalization”).

Like the contrast energy model, the untuned normalization model predicts a similar BOLD amplitude for snakes and gratings, thereby failing to account for the observation about the two stimulus classes (**Figure 5, upper panel**). The model is not sensitive to heterogeneity across orientations because the normalization pool equally weights all orientation channels. Therefore, the output does not depend on whether the contrast energy is concentrated in one orientation channel as in the gratings or spread across many channels as in the snakes. Like the contrast energy model, the untuned normalization model’s failure to account for the greater response to snakes is confirmed by low variance explained in each of the four data sets with these target stimuli (**Figure 8**, **Figure S2b**).

**Figure 5.**
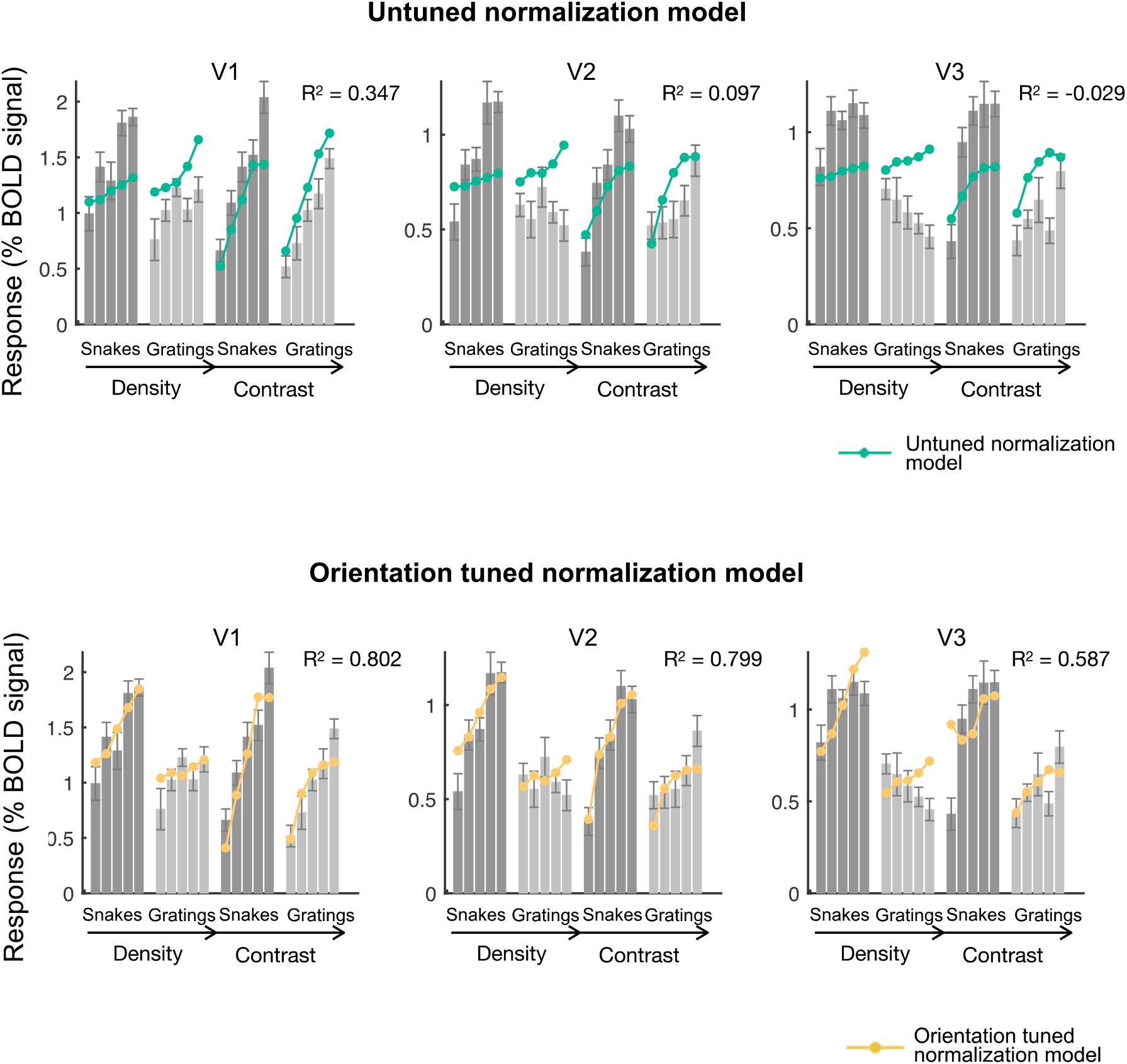
The tuned normalization model is much more accurate than the untuned normalization model. The mean fMRI responses from V1, V2, and V3 are replotted from Figure 3. The dots are the cross-validated predictions from the untuned normalization model (above) or the tuned normalization model (below). Data and model fits plotted from the function *s4_visualize(’figure5A’) and s4_visualize(’figure5B’)* in the <u>code repository</u>. See *Figure S2b-c* for fits to all 4 datasets.

### 2.4. The larger response to snakes is captured by an orientation tuned normalization model

The untuned normalization model implements a surround suppression that includes energy at all orientations. Findings from electrophysiology (<u>Cavanaugh, Bair, and Movshon 2002</u>; <u>DeAngelis,</u> <u>Freeman, and Ohzawa 1994</u>; <u>Sillito et al. 1995</u>; <u>Shushruth et al. 2013</u>; <u>Trott and Born 2015</u>), psychophysics (<u>Xing and Heeger 2001</u>; <u>Wang, Heeger, and Landy 2012</u>; <u>Shushruth et al. 2013</u>; <u>Solomon,</u> <u>Sperling, and Chubb 1993</u>), neuroimaging (<u>Bloem and Ling 2019</u>; <u>Klimova, Bloem, and Ling 2021</u>; <u>Zenger-</u> <u>Landolt and Heeger 2003</u>; <u>McDonald et al. 2009</u>; <u>Joo, Boynton, and Murray 2012</u>), and theory (<u>Schwartz</u> <u>and Simoncelli 2001</u>), however, suggest that surround suppression is orientation-tuned. For example, Cavanaugh, Bair, and Movshon (<u>2002</u>) reported that the response of a neuron to a stimulus at its preferred orientation in its receptive field is suppressed more when the surrounding region contains contrast at the same orientation compared to different orientations. Because our grating stimuli have contrast energy concentrated at a single orientation, and the snake stimuli do not, one might surmise that an *orientation tuned normalization* model would show greater suppression for the gratings, where the RF centers and surrounds will have matched orientations, than for the snakes, where the orientations are more likely to differ between center and surround. If so, this could then account for our observed effect.

The untuned normalization model is the same as the tuned model except for one difference: the tuned model normalizes the contrast energy across nearby locations only at the preferred orientation (orientation-tuned surround) (**Figure 4**, lower right panel, diagonals). Within a single location, the normalization is untuned (off-diagonals in the same panel), also called cross-orientation suppression (<u>Morrone, Burr, and Maffei 1982</u>).

The orientation tuned normalization model captures the large and systematic differences in response amplitude between gratings and snakes (**Figure 5**, **lower panel**). In the example dataset, the BOLD amplitude and the orientation tuned normalization model predictions are about a half for the gratings compared to the snakes, both for the density and contrast stimulus manipulations, and for all three visual areas (**Figure 7**). In addition to capturing this difference in the means between the two stimulus classes, the model also captures the difference in slope. As the density or contrast increases, the model predicts steeper slopes for snakes than gratings. These patterns, both in the data and in the model fits, were found across the four data sets (**Figure S2c**). The orientation tuned normalization model is more accurate than the previous two models in all cases (**Figure 8**, four datasets and three ROIs). The pattern also held up for different size surrounds – what mattered was whether the surround was tuned or not tuned, not its size (data not shown). The indifference to the size of the surround almost certainly reflects the properties of the stimulus set, not neural tuning: the stimuli are all textures, with similar properties across the image. Had the stimuli varied systematically across location, the size of the surround would likely have had a large effect on model accuracy.

### 2.5. Normalization by orientation anisotropy

The large advantage in prediction accuracy of the tuned over the untuned normalization model supports the idea that suppression is feature specific. As with any model, we chose a specific instantiation of a more general idea, here feature-specific suppression. The specific instantiation entailed a minimal change from the untuned normalization model, requiring only a change in normalization weights, and builds on the tradition of feedforward, filter-based models. Feature-specific tuning can also be implemented in other ways, for example based on more abstract ideas like predictability or redundancy in the image. There is some evidence in favor of models like these (<u>Coen-Cagli, Kohn, and Schwartz</u> <u>2015</u>; <u>Vinck and Bosman 2016</u>). We implemented a second method of achieving orientation-tuned normalization, in which normalization was proportional to *orientation anisotropy* (**Figure 7**, Normalization by orientation anisotropy, “NOA”). Normalization in this model is most pronounced when the surround and center orientations match, not necessarily when the surround orientation matches the *preferred tuning* of the center (See *3.3 What is the tuning in orientation-tuned normalization?*).

Specifically, in the normalization by anisotropy model the contrast energy is normalized by the standard deviation across the outputs of the orientation channels (**Figure 7**). This normalization by anisotropy model applies greater normalization when the contrast energy is concentrated in a single orientation channel, resulting in a lower response for gratings. This implementation is consistent with the idea that responses are reduced by the amount of redundancy in the image.

The normalization by anisotropy model exhibits similar predictions to the orientation-tuned normalization model, capturing the larger response to the snakes (**Figure 6**). Both models predict that for these stimuli, the responses to snakes are about double the responses to gratings, similar to the data (**Figure 7**). It also predicts a steeper slope for the snakes than the gratings, both as a function of density and contrast. The successes of these models validate the idea that normalization depends on how contrast energy is distributed across orientations, not just on the overall contrast energy.

**Figure 6.**
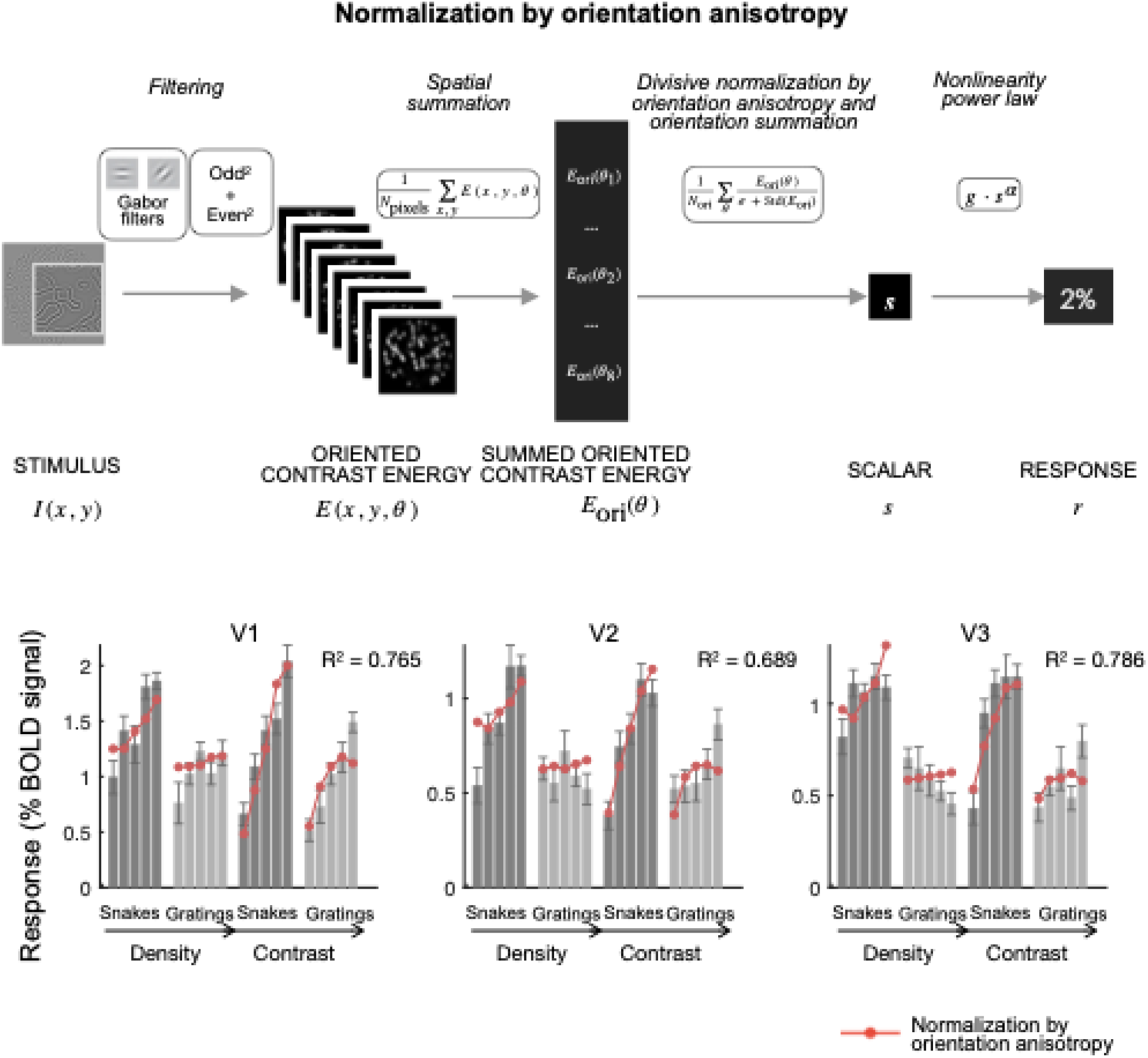
The normalization by orientation anisotropy model also accounts for the responses in V1-V3. The mean fMRI responses from V1, V2, and V3 are replotted from Figure 4. The red dots are the cross-validated predictions from the normalization by orientation anisotropy model. Data and model fits plotted from the function *s4_visualize(’figure6’)* in the <u>code repository</u>. See *Figure S2d* for fits to all 4 datasets.

**Figure 7.**
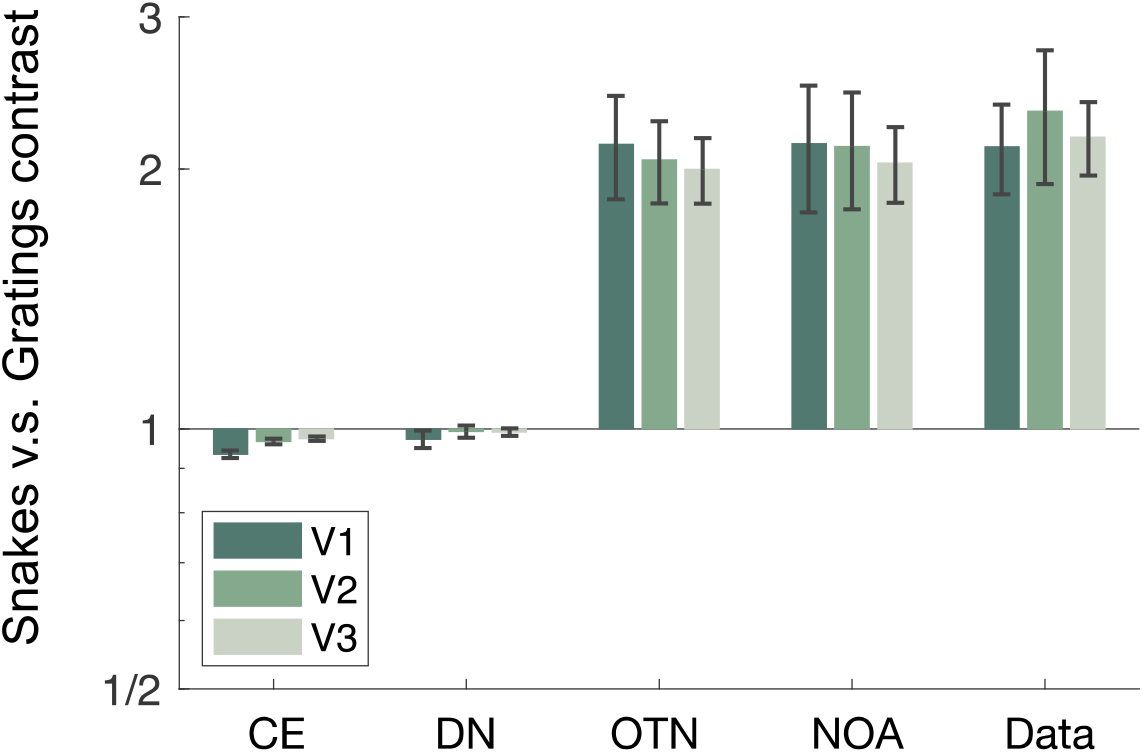
The orientation dependent models account for the higher responses to snakes than gratings. For each visual area, V1-V3, we averaged the response to snakes and to gratings across stimuli in the target set and across the 4 data sets to compute the ratio of snakes to gratings: mean(snakes) / mean(gratings). We computed this value for each dataset and plotted the average and standard error across the four datasets. In the data, the response to snakes is about double to gratings. This is matched in the orientation-tuned normalization model (OTN) and normalization by anisotropy model (NOA) predictions but not the predictions from other models (contrast energy (CE), and untuned normalization (DN)). Data and model fits plotted from the function *s4_visualize*(’figure7’) in the <u>code repository</u>.

**Figure 8.**
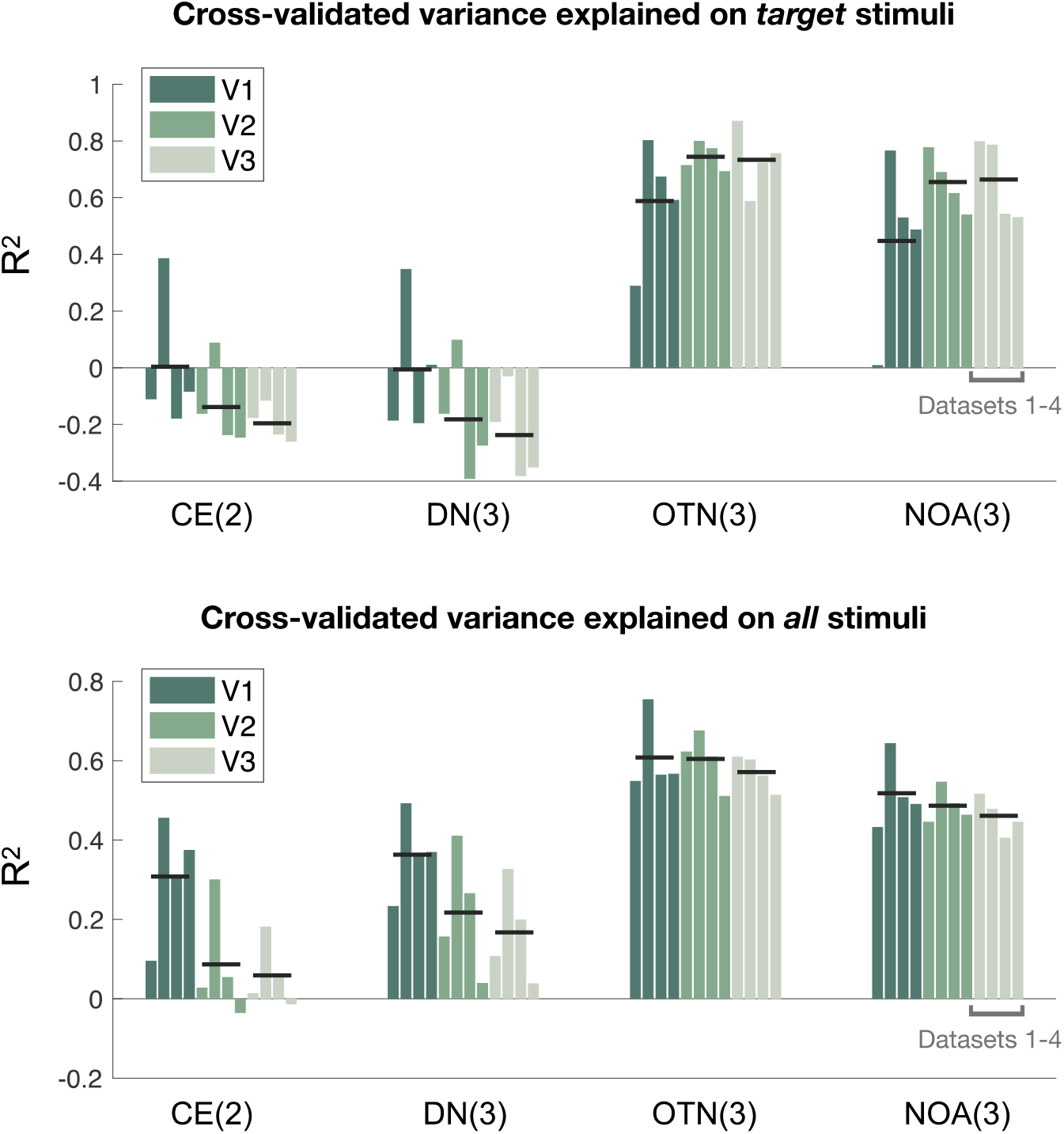
Cross-validated variance explained for 4 models across all ROIs and data sets. The figure shows the leave-one-out cross validated coefficient of determination (R^2^) for four data sets in three visual areas and four models. The number of fitted model parameters (degrees of freedom) is indicated in parentheses for each model type. The upper panel contains the R^2^ for the target data set, comprising 18 stimuli for data set 1 (DS1) and data set 2 (DS2) and 17 for data set 3 (DS3), data set (DS4). The lower panel shows the R^2^ for a larger set of data (50 for data set 1; 48 for data set 2; 39 for data set 3 and 4). The four bars in each group correspond to datasets 1-4. The black horizontal lines are the means across the 4 datasets. The R^2^ values are also reported in Supplementary table S4. Data plotted from the function *s4_visualize*(’figure8’) in the <u>code repository</u>.

### 2.6. The two normalization models that are sensitive to orientation accurately predict responses to a wide range of stimuli

The good performances of the tuned normalization model and the anisotropy model demonstrate that sensitivity to orientation is an important part of normalization. This sensitivity is needed to capture the larger response to the gratings than the snakes. Overall, across 4 models, accuracy clusters into two groups: low accuracy for the two models without orientation sensitivity and high accuracy for the two models with orientation sensitivity (**Figure 8**, upper panel). The accuracy differences across models are not due to the number of free parameters: the untuned normalization model has the same number of free parameters (three) as the tuned normalization model and the anisotropy model. Moreover, the prediction accuracy was summarized by *n*-fold cross-validation, so that more parameters do not necessarily lead to more accurate predictions. The advantage for the two models with orientation sensitivity dwarfs the slight difference between these two models, although we note that the tuned normalization model has a numerically higher accuracy than the anisotropy model for nearly all data sets in all conditions (**Table S4**). Interestingly, for the two untuned models, prediction accuracy declines from V1 to V2 to V3, whereas for the two tuned models, accuracy increases from V1 to V2/V3 (black lines in **Figure 8**).

Because the motivation for implementing the two orientation sensitive models was, in part, to explain the greater response to snakes than gratings, it is important to test the models for other stimuli as well. We refit all 4 models to the full data sets shown to our four subjects, which consisted of 50 (data set 1), 48 (data set 2) and 39 (data set 3, data set 4) stimuli, spanning a variety of texture types. In addition to the snakes and gratings, there are textures we refer to as *noise bars*, *waves*, *plaids*, and *circular* (**Supplementary Figure 1** and **Table S3**).

Just as with the target stimuli, across the full sets of stimuli, the tuned normalization model and the anisotropy model made accurate predictions, explaining 60%-76% and 48%-64% of the cross-validated variance in V1-V3 for the example data set (**Figure 9**). These two models also provide good fits to the other three datasets, shown in Figures S3-S6. The fits to the larger stimulus sets, like the fits to the target stimuli alone, capture the observation about the two stimulus classes, meaning a larger predicted response for snakes than gratings. The two models also accurately predict lower responses to waves (one dominant orientation) than noise bars (many orientations). The two models also predict increasing response amplitudes from gratings (one orientation) to plaids (two orientations) to circular (16 orientations), as evident in stimulus sets 3 and 4 (**Supplementary figures 5cd, 6cd**). This pattern of predictions matches the data. The untuned models do not differ in their predictions for these three stimulus categories.

**Figure 9.**
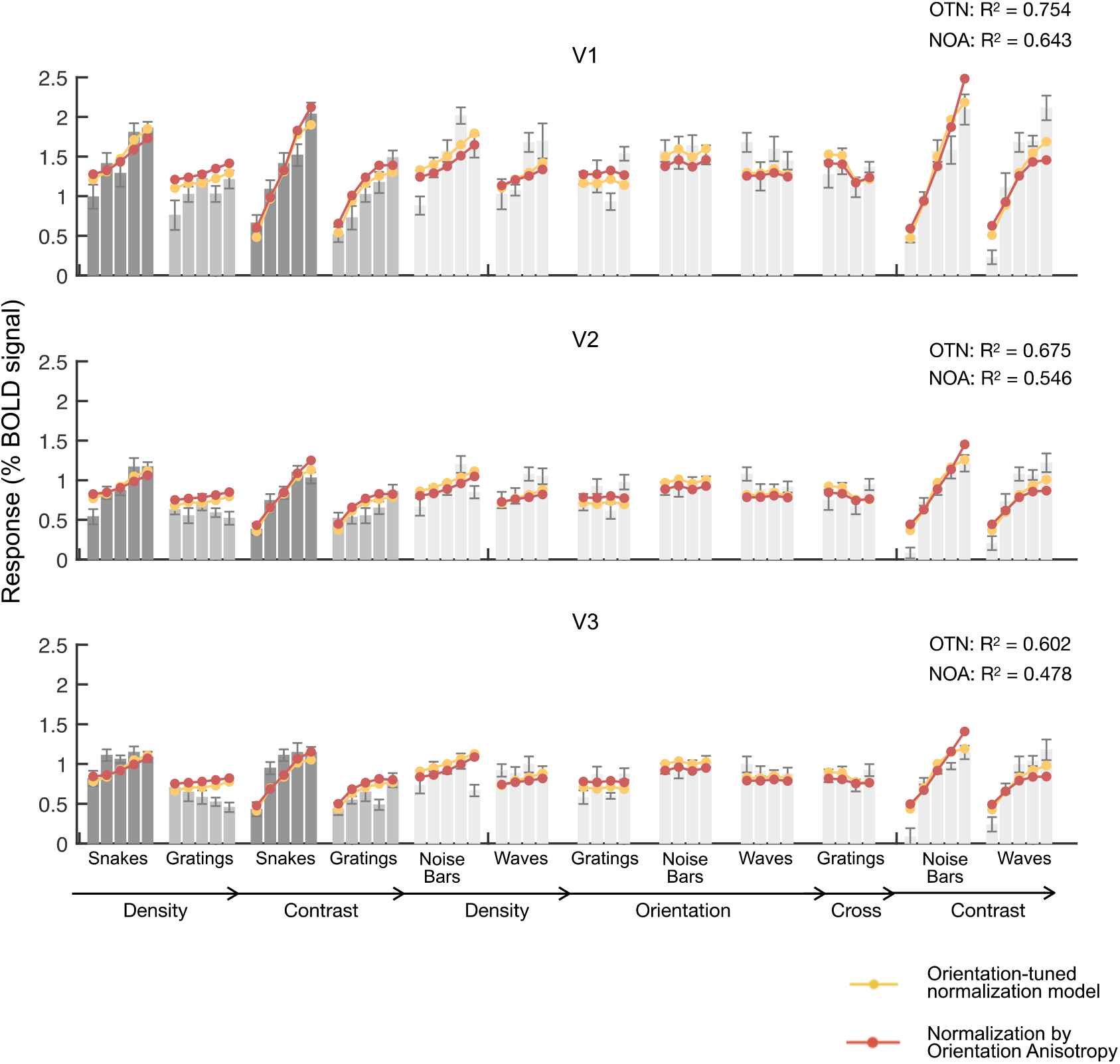
The orientation dependent models fit a wide variety of stimuli. The mean fMRI responses from V1, V2, and V3 are shown for the full set of stimuli from data set 2 (See Supplementary Figure S1 for images). The data for snakes and gratings are replotted from figures 3, 5-6, and again shown in dark and light gray. An even lighter gray is used for all other stimulus classes. The red and yellow dots are the cross-validated predictions from the orientation-tuned normalization model and the normalization by orientation anisotropy model. See Supplementary Figures S3-S6 for similar plots for data sets 2-4. Data and model fits plotted from the function *s4_visualize*(’figure9’) in the <u>code repository</u>.

Across ROIs and data sets, the orientation tuned normalization model accounts for the highest variance, with R^2^ ranging from 51% to 76% (**Figure 8**, bottom; **Table S4**, the third row). The anisotropy model ranks second in all cases, substantially outperforming the two baseline models. Similar to the pattern with the target stimuli, when fitting to all stimuli the two untuned models show substantially decreasing accuracy from V1 to V2 to V3. The tuned normalization model and the anisotropy model decrease only slightly in accuracy from V1 to V2 to V3, meaning that the advantage for the tuned models is largest in extrastriate areas.

## 3. Discussion

### 3.1 Why do some models fail to account for the much larger response to snake stimuli?

We began with the observation that the BOLD response is about twice as large for patterns with curved contours (snakes) as for similar stimuli with straight, parallel contours (gratings). The difference in the responses is not a peculiarity of the BOLD signal, as a similar pattern was also observed in human intracranial measures of the field potential (broadband power from ECoG electrodes). The contrast energy model without normalization and the contrast energy model with untuned normalization did not predict large differences between the responses to the two stimulus classes. Both models pool contrast energy over space and orientation without an orientation-specific normalization or other orientation specific non-linearity. If two images have the same total contrast energy, then the way the energy is distributed across orientation channels will not matter, either for the measure of total energy (contrast energy model) or for the amount of normalization (untuned normalization model).

The implementation of the orientation-tuned normalization model was motivated in part from electrophysiology results showing that in V1, surround suppressive fields tend to be tuned to orientations close to the RF center’s preferred orientation (<u>Cavanaugh, Bair, and Movshon 2002</u>). Psychophysical experiments also show interactions between a target and surround that depend on matched orientation (<u>e.g., Solomon, Sperling, and Chubb 1993</u>). While there is a lot of evidence from electrophysiology, fMRI, and psychophysics in support of feature tuning in normalization, to our knowledge this is the first image-computable model of orientation-tuned normalization fit to neural data from a wide range of images. Implementing the model and fitting it to data enabled us to assess whether (1) it quantitatively accounts for two-fold difference in response to snakes vs gratings (2) whether it also provides good fits to a wide range of other stimuli that it was not explicitly implemented for and (3) whether it can capture other more subtle effects, like the difference in the slope of the contrast response function between snakes and gratings. In all three cases, the answer was yes. We also implemented a second model with tuned normalization (the normalization by orientation anisotropy model), and this model also provided an affirmative answer to these questions, confirming the importance of including orientation dependence in the normalization computation. We discuss similarities (3.2) and differences (3.3) between the two models below.

### 3.2 Model behavior: Models with orientation dependent normalization capture the differences in both *mean* and in *slope* between snakes and gratings

The two models with orientation dependent normalization predict larger outputs, *on average*, for snakes than gratings, even when the contrast energy of the stimuli is approximately matched. This is due to how the normalization is computed. The grating stimuli elicit large outputs in the orientation channels that are matched to the stimulus, moderate outputs in adjacent orientation channels, and little to no response in other channels. As a result, there is high anisotropy (standard deviation across channel outputs), resulting in more suppression in the normalization by orientation anisotropy model. There is also high suppression from the surround in the orientation-tuned normalization model. The two models were implemented to capture the difference in mean between the two stimulus classes, so perhaps it is not so surprising that they do so.

The two models, unlike the contrast energy model and the untuned normalization model, also accurately predict *steeper slopes* for snakes than gratings (with respect to both contrast and density), a pattern that the models were not explicitly motivated to capture. They predict the difference in slope because at low total contrast energy for the image, there is little normalization, and hence the response to a snake and a grating stimulus will be comparable. This explanation applies to both low contrast stimuli and low-density stimuli, because in both cases the summed contrast energy is low, meaning the normalization term in the denominator is small. Hence, little normalization at low contrast is expected from the model, and is also confirmed by empirical measures of spatial summation and surround suppression at different contrast levels {Sceniak, 1999 #3178}{Cavanaugh, 2002 #899}. The more nuanced predictions from these models is that for stimuli with high total contrast energy (high stimulus contrast and high density), there is a lot of normalization for gratings and much less for snakes, resulting in a more pronounced difference in predicted response. The difference in predicted responses at high contrast but not at low contrast causes a difference in slope. For the two untuned models, normalization increases with contrast energy, but it increases similarly for the snakes and gratings, hence predicting similar slopes.

Interestingly, the data and the two models with orientation dependent normalization also show a greater slope for “noise bar” stimuli than “waves” (Supplementary figures S3cd, S4cd). The noise bars, like the snakes, have many orientations in first order contrast (but unlike the snakes, have only one dominant orientation for second order contrast). The waves are the complement, with one dominant orientation for first order contrast (like gratings) but many orientations for second order contrast. The pattern in the data and model predictions is that the stimuli with many orientations (noise bars) increase more steeply as a function of contrast than the stimuli with a narrower range of orientations (waves), supporting the observations made with snakes and gratings, but adding the further nuance that what seems to matter most is the orientation distribution (wide vs narrow) for first order rather than second order contrast.

The difference in slopes for snakes vs gratings, both in the data and model predictions, is related to, but not identical to, results observed for single unit V1 cells. For a typical V1 cell, the contrast response function has a higher slope for preferred than for non-preferred stimuli (<u>Li and Creutzfeldt 1984</u>; <u>Albrecht and Hamilton 1982</u>; <u>Sclar and Freeman 1982</u>), a pattern also predicted by normalization models (<u>Heeger 1992b</u>). The pattern is predicted because of the difference in the *numerator*, which is high for preferred stimuli and low for non-preferred. The denominator is about the same for the two stimuli, proportional to the total contrast energy in the stimulus. We attribute the difference in slope between snakes vs gratings to a difference in the *denominator* of the normalization equation, which is high for gratings, low for snakes. The numerator is about the same for the two classes, proportional to the total contrast energy. In both cases–single unit responses to non-preferred stimuli, and population responses to grating stimuli– the reduced slope is due to a large amount of normalization relative to the driven response.

### 3.3 What is the tuning in orientation-tuned normalization?

We implemented two models with orientation dependent normalization. Both capture the same tendency for more normalization for homogeneous stimuli like gratings. They differ in implementation, however. The tuned normalization model differs only slightly from more typical normalization models: the only difference is that the normalization weights happen to conform to a specific pattern, such that cells with similar feature tuning (here, orientation) with nearby receptive fields have high weights, and cells with different feature tuning have low weights. As a result, surround suppression is most effective according to this model when the orientation of a surrounding region is matched to the preferred orientation of a cell. This is justified by findings from single unit data in macaque V1 that “the surround influence was always suppressive when the surround grating was at the neuron’s preferred orientation” (<u>Cavanaugh, Bair, and Movshon 2002</u>).

The normalization by orientation anisotropy model is a larger departure from the standard normalization model, since the computation of anisotropy is not a simple weighting of the outputs of nearby cells. Unlike the tuned normalization model, it has the greatest suppressive effect when the features of a stimulus are anisotropic (like a grating) irrespective of whether the stimulus orientation matches the center tuning of a cell. Interestingly, there is also empirical support for this pattern from the same study by Cavanaugh et al (<u>2002</u>): specifically, evidence for “the tuning of the surround being dependent to some degree on the stimulus used in the center—suppression was often stronger for a given center stimulus when the parameters of the surround grating matched the parameters of the center grating even when the center grating was not itself of the optimal direction or orientation.” This pattern has been found in multiple studies, indicating that for macaque and cat V1 cells, surround suppression is often maximal when the surround and center stimulus orientation match, independent of the orientation preference of the cell (<u>Sillito et al. 1995</u>; <u>Shushruth et al. 2012</u>; <u>Angelucci et al. 2017</u>).

Put another way, the tuning of the surround changes as the center orientation or direction changes. As shown by simulation, our normalization by anisotropy model can capture this observation from single units, but our tuned normalization model cannot (**Figure 10**).

**Figure 10.**
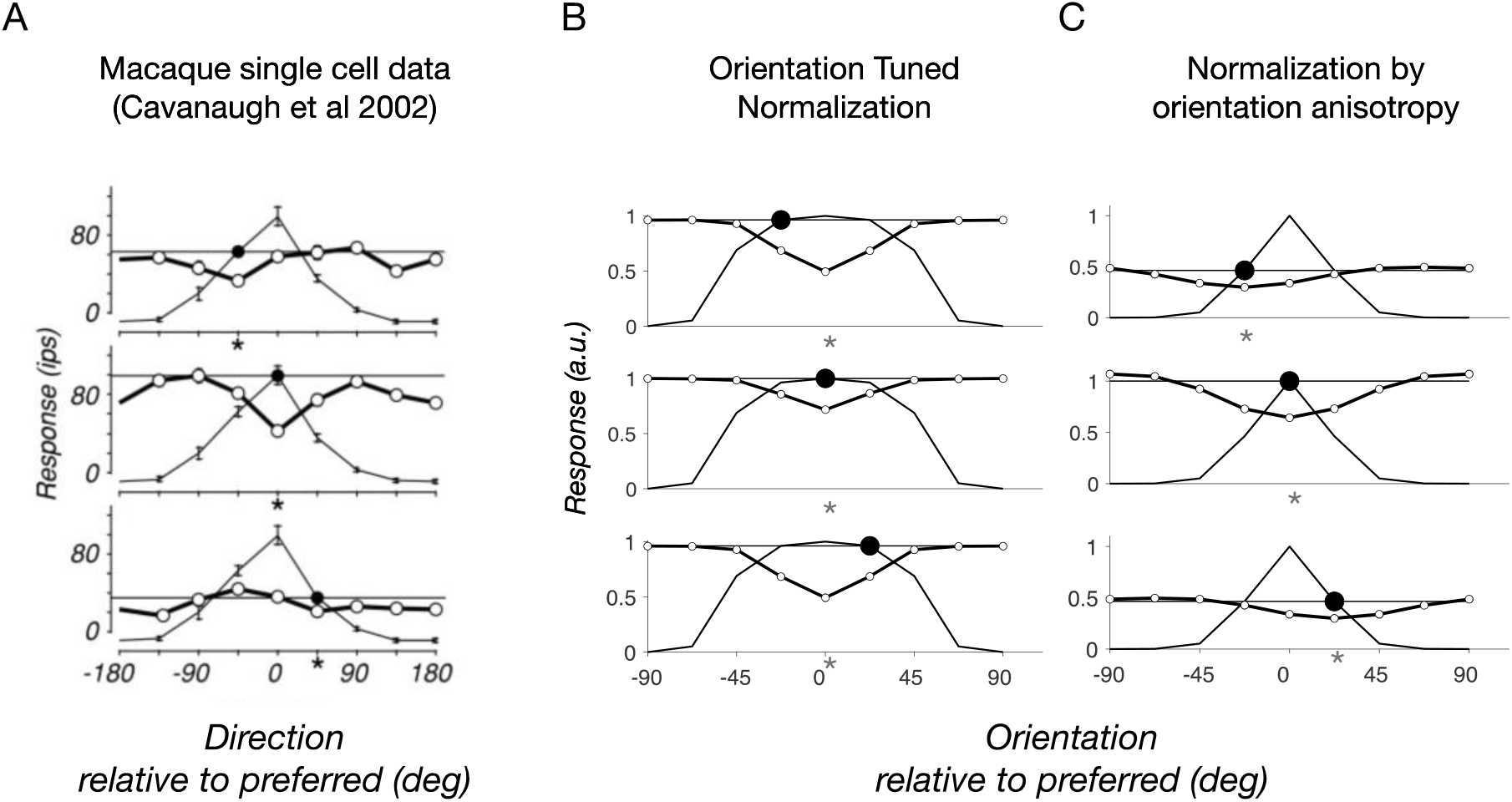
Center surround interactions in single units and computational models. (A) Data replotted from three example cells from macaque V1 (<u>Cavanaugh, Bair, and Movshon</u>). The thin lines are tuning curves for drifting gratings in the cell’s RF center, with 0 deg indicating the preferred direction. The thick line and open circles show responses to stimuli including a center and surround. The center direction is indicated by the large black dot, and the surround direction is indicated by the values in the x-axis. The important observation is that the greatest suppression (indicated by the asterisks) occurs when the surround direction matches the center direction, not when the surround direction matches the center’s preferred direction. (B) Simulations for the orientation-tuned normalization model for stimuli similar to those in panel A (but static rather than drifting). Because the surround suppression is matched to the center’s preferred tuning, the largest suppression occurs at 0 deg, differing from the single unit data. (C) Simulations for the normalization by orientation anisotropy model for the same stimuli as in panel B. Here, the greatest suppression is when the surround orientation matches the center orientation, similar to the single unit data. Data and simulations plotted from the function *s4_visualize*(’figure10’) in the <u>code</u> repository.

The two models we implemented leave us in an unusual position. We have one model that has slightly higher prediction accuracy and is more in line with standard normalization models (orientation-tuned normalization), and a second model with slightly lower (but still high) prediction accuracy, but a closer fit to some data from single units (normalization by orientation anisotropy). The simplest conclusion is that a contrast energy model incorporating some form of orientation-dependent normalization greatly outperforms models that do not have tuned normalization. The specific circuit-level implementation of orientation-dependent normalization is not sufficiently constrained to clearly favor one or the other of these two computations. Moreover, given the diversity of cells in visual cortex, it is not likely that a single, simplified model will be sufficient to capture the behavior of all cells or cell populations for all stimuli. A neural network model with recurrence, temporal dynamics, and with feedforward, feedback and lateral connections might provide insight into the specific ways in which the surround modulation emerges (Shushruth et al. 2012; <u>Heeger and Zemlianova 2020</u>).

Interestingly, the denominator in the normalization by anisotropy model (the normalizer) is almost the same as the numerator of an orientation variance model used to predict the amplitude of gamma oscillations measured with ECoG electrodes in human subjects (Hermes et al., 2019). The fact that the same term is used in the numerator to predict gamma oscillations and the denominator to predict BOLD is consistent with the empirical observation that many of the stimuli that are most effective for driving large gamma oscillations (high contrast luminance gratings) are relatively ineffective for eliciting BOLD signals (<u>Butler et al. 2017</u>; <u>Hermes, Nguyen, and Winawer 2017</u>), and multiunit action potentials (<u>Hermes, Kasteleijn-Nolst Trenite, and Winawer 2017</u>; <u>Ray and Maunsell 2011</u>). The link may be that gamma oscillations, rather than being the fundamental mechanism for perception and long-range cortical communication (<u>Engel and Singer 2001</u>; <u>Fries 2005</u>), are rather a result of the normalization process (<u>Hermes et al. 2019</u>; <u>Ray, Ni, and Maunsell 2013</u>). An open question is whether gamma oscillations might be predicted as accurately, or perhaps more accurately, by the normalization pool used in our orientation-tuned normalization model than the normalization pool of the normalization by anisotropy model.

### 3.4 Orientation dependent normalization and image statistics

The normalization by orientation anisotropy model and the orientation-tuned model showed larger advantages over untuned models for V2 and V3 compared to V1. This pattern is consistent with findings from electrophysiology and computational modeling of behavior. First, evidence from the non-human primate visual system suggests that the orientation tuning of surround suppression in V1 arises in large part from feedback from extrastriate areas, especially for the more spatially distant effects of surround suppression (<u>Angelucci et al. 2002</u>; <u>Angelucci et al. 2017</u>; <u>Shmuel et al. 2005</u>). If the tuned suppressive effects depend on computations in extrastriate regions, then these regions may also exhibit more tuned suppression than V1.

More generally, tuned suppression, either in the form of our tuned normalization model or the anisotropy model, reflects sensitivity to higher-order image statistics (correlations over space between filter outputs). Models of extrastriate neural responses, especially V2, also tend to be more sensitive to higher-order image statistics than to contrast energy *per se*, such as models based explicitly on texture statistics (<u>Freeman and Simoncelli 2011</u>; <u>Freeman et al. 2013</u>; <u>Movshon and Simoncelli 2015</u>; <u>Okazawa,</u> <u>Tajima, and Komatsu 2017</u>){Ziemba, 2021 #3187}, or models with multiple subunits that may exhibit heterogenous feature tuning {Liu, 2016 #3184}{Willmore, 2010 #3186}{Tao, 2012 #3185}.

We consider one specific way that our normalization by anisotropy model might be linked to the V2 models proposed by Simoncelli and colleagues. Suppose the V1 population computes contrast energy localized in orientation and space. (For simplicity, we ignore spatial frequency, as we did here experimentally by band-pass filtering our stimuli.) We then suppose that the V2 cells compute various weighted sums of the V1 outputs. Specifically, we assume that, for each spatial location, the weights among V2 cells form a Fourier basis set on the V1 outputs (across orientation). If these weights are arranged in pairs (similar to the odd and even V1 filters), and the outputs are squared and summed across phase (similar to the V1 energy model), then then the summed V2 population output will be proportional to the variance in the V1 response across orientation (David Heeger, personal communication). If this population output is the normalization pool for V2, then we get a model like the anisotropy model, which normalizes by the standard deviation across orientation channels.

### 3.4 Are typical contrast energy V1 models missing something important?

No model is complete. The standard normalized contrast energy model of V1, when fit with appropriate parameters, can capture substantial variance in V1 responses (<u>Carandini, Heeger, and Movshon 1997</u>), but not all. Our results indicate that the model fails for at least some relatively simple stimulus classes, and that the failure can be large. But whether a more complex model is needed, such as a gated normalization model (<u>Coen-Cagli, Kohn, and Schwartz 2015</u>) or either of the orientation dependent normalization models we implemented, will depend on the stimulus set tested and the purpose of modeling. It is reasonably likely that the computations in both the anisotropy and the tuned normalization models are more connected to computations in extrastriate visual maps than in V1, but may also influence the response in V1 via feedback. (<u>Angelucci et al. 2017</u>; <u>Henry et al. 2020</u>).

More generally, since the development of divisive normalization models in the early 1990s, there has not yet been a generally agreed up description of exactly what cell populations (and with what weights) contribute to normalization. Some attempts have been made from assuming efficient coding of natural images (<u>Schwartz and Simoncelli 2001</u>) or from fitting model parameters to neural data (<u>Burg et al.</u> <u>2021</u>). In this sense, a standard model of V1, with parameters set, that is downloadable and executable on arbitrary input images, does not yet really exist.

Both the anisotropy model and the tuned normalization model we presented make some advance but also have some important limits, most notably the lack of spatial receptive fields, as well as a lack of sensitivity to second-order contrast. Hence, they do not supersede other models, but rather provide compact computational summaries of response patterns that are not well captured by other models. Integrating better computational tools for validation (<u>Lerma-Usabiaga et al. 2020</u>), model-based stimulus development (<u>Golan, Raju, and Kriegeskorte 2020</u>; <u>Wang and Simoncelli 2008</u>), high-quality standardized datasets (<u>Allen et al. 2021</u>), and theory (<u>Simoncelli and Olshausen 2001</u>), may offer the best path toward more complete understanding of neural circuits in visual cortex.

## 4. Methods

We analyzed and modeled four fMRI data sets for this paper. Data sets 1 and 2 were collected at NYU. Data sets 3 and 4 are re-analyzed from a previous paper (<u>Kay, Winawer, Rokem, et al. 2013</u>), for which the fMRI data and stimuli are freely available online (http://kendrickkay.net/socmodel/).

### 4.1 Participants

The two NYU participants were both experienced MRI subjects (female; 24, 29 yo). Data were collected at NYU’s Center for Brain Imaging. The experimental protocol was approved by the University Committee on Activities Involving Human Subjects, and informed written consent was obtained from the participants before the study. Both participants had corrected-to-normal vision. The subjects participated in two separate scanning sessions, one for retinotopic mapping and one for the main study on encoding of textures.

### 4.2 Stimuli

Publicly accessible links to the stimuli, the names of the stimulus classes, and the correspondence between our naming convention and those in the Kay *et al*. (<u>2013</u>) paper are described in Supplementary Tables 1 and 2.

#### 4.2.1 Stimuli for data sets 3 and 4

Data sets 3 and 4 correspond to subject 1 and subject 2, respectively, in Kay *et al*. (<u>2013</u>). The stimuli for the two subjects were the same and are referred to as “Stimulus set 2” on the website (http://kendrickkay.net/socmodel/). The publicly available stimulus set includes 156 stimuli, 39 of which were used for this paper (**Supplementary Tables 2 and 3**). The reason we use only a subset of the stimuli is that we modeled how visual areas respond to textures ignoring retinotopic preference, and many of the stimuli used in the original paper varied systematically over space in order to map spatial receptive fields. We used only the large-field textures, i.e., the subset of stimuli whose patterns were similar across the whole circular aperture.

The 39 stimuli are organized into 7 groups, each of which we describe with two terms, one term for the type of texture and one term for the way in which the stimuli within the group vary. For example, GRATINGS (contrast) are stimuli which come from the grating family and vary from low to high contrast. GRATINGS (density) stimuli come from the same family but have uniform contrast and vary in the spacing between the contours. The correspondence between how we refer to the stimuli and how Kay et al. referred to them is in **Supplementary Table 2**. Below we describe the general stimulus characteristics and the 7 specific classes used for this paper. Most of the text is duplicated from Kay *et al*. (<u>2013</u>) (p. 11), indicated by italics.

General stimulus characteristics. Stimuli were constructed at a resolution of 256 pixels x 256 pixels and were upsampled to 800 pixels x 800 pixels for display purposes. All stimuli were presented within a circular aperture filling the height of the display; the rest of the display was filled with neutral gray. The outer 0.5 deg of the circular aperture was smoothly blended into the background using a half-cosine function.

Stimuli consisted of grayscale images restricted to a band-pass range of spatial frequencies centered at 3 cycles per degree. To enforce this restriction, a custom band-pass filter was used in the generation of some of the stimuli. The filter was a zero-mean isotropic 2D Difference-of-Gaussians filter whose amplitude spectrum peaks at 3 cycles per degree and drops to half-maximum at 1.4 and 4.7 cycles per degree. Restricting the spatial frequency content of the stimuli avoids the complications of building multiscale models and helps constrain the scope of the modeling endeavor. Even with the spatial frequency restriction, it is possible to construct a rich diversity of stimuli including objects and other naturalistic stimuli…. Each stimulus consisted of nine distinct images that were presented in quick succession. The purpose of this design was to take advantage of the slow dynamics of the BOLD response and average over stimulus dimensions of no interest (e.g., using sinusoidal gratings differing in phase to average over phase).

A key motivating observation for this paper is that the response to gratings was lower than to curved stimuli. Four groups of stimuli, two groups of gratings and two groups of snakes, were studied first and are referred to throughout the paper as *target stimuli.* These are described first.

### Target stimuli

SNAKES (contrast, 10 stimuli). Kay et al. refers to these stimuli as noise patterns: Noise patterns were created by low-pass filtering white noise at a cutoff frequency of 0.5 cycles per degree, thresholding the result, performing edge detection using derivative filters, inverting image polarity such that edges are black, and applying the custom band-pass filter (described previously). We generated nine distinct noise patterns and scaled the contrast of the patterns to fill the full luminance range. [The contrast stimuli were then] constructed by varying the contrast of the noise patterns. Ten different contrast levels were used: 1%, 2%, 3%, 4%, 6%, 9%, 14%, 21%, 32%, and 50%. These contrast levels are relative to the contrast of the patterns used in SPACE [not used in this study], which is taken to be 100%.

SNAKES (density, 5 stimuli). These stimuli used the same type of noise patterns as SPACE [not used here] but varied the amount of separation between contours. We generated noise patterns using cutoff frequencies of 2.8, 1.6, 0.9, 0.5, and 0.3 cycles per degree, and numbered these from 1 (smallest separation) to 5 (largest separation). The noise patterns used in SPACE correspond to separation 4; thus, we only constructed stimuli for the remaining separations 1, 2, 3, and 5. The noise patterns occupied the full stimulus extent (no aperture masking) GRATINGS (contrast, 4 stimuli). These stimuli consisted of horizontal sinusoidal gratings at 2%, 4%, 9%, and 20% Michelson contrast. The spatial frequency of the gratings was fixed at 3 cycles per degree.

GRATINGS (density, 5 stimuli). The highest density stimulus in this group is similar to the horizontally oriented stimulus in GRATINGS (orientation), i.e., similar to a horizontal high-contrast grating, but it is not precisely a sinusoidal grating. It is made by convolving equally spaced horizontal lines with the custom band-pass filter (described previously). When the gratings are spaced appropriately (⅓ deg spacing) and filtered by a band-pass filter centered at 3 cycles per deg, the result is close to a sinusoidal grating at 3 cycles per deg. When the spacing is larger, there are several parallel band-pass contours with uniform gray between them. The spacing between parallel lines for the 5 stimuli varied in powers of 2, as 1/3 deg x 1,2,4,8, or 16, from densest to sparsest. Because the grating and snakes stimuli were both constructed by convolving lines with the same band-pass filter, they have some similar properties. They differ in that the lines here were straight whereas the lines used for constructing the snakes stimuli were curved.

### Additional stimuli

GRATINGS (orientation, 8 stimuli). These stimuli consisted of full-contrast sinusoidal gratings at eight different orientations. The spatial frequency of the gratings was fixed at 3 cycles per degree. Each [of the 9 exemplars per stimulus] consisted of gratings with the same orientation but nine different phases (equally spaced from 0 to 2π).

PLAID (contrast, 4 stimuli). These stimuli consisted of plaids at 2%, 4%, 9%, and 20% contrast (defined below). Each condition comprised nine plaids, and each plaid was constructed as the sum of a horizontal and a vertical sinusoidal grating (spatial frequency 3 cycles per degree, random phase). The plaids were scaled in contrast to match the root-mean-square (RMS) contrast of the GRATING stimuli. For example, the plaids in the 9% condition were scaled such that the average RMS contrast of the plaids is identical to the average RMS contrast of the gratings in the 9% GRATING stimulus.

CIRCULAR (contrast, 4 stimuli). These stimuli were identical to the PLAID stimuli except that sixteen different orientations were used instead of two.

#### 4.2.2 Stimuli for data set 1

Data set 1 was collected at NYU. The data set was designed to replicate some of the effects observed from data sets 3 and 4 (the greater response to snakes than gratings), but also to extend the measurements to new stimulus classes. The general stimulus characteristics were the same as those used in data sets 3 and 4. However, because the display size differed, the image resolution in pixels also differed (400 x 400 here, vs 800 x 800 above), and there were slight differences in the bandpass filter. The stimulus size in degrees of visual angle was the same (12.5 deg diameter). A total of 50 stimuli were tested. (The numbers below total more than 50 because some stimuli belong to more than one group, as indicated in **Supplementary Table 2**).

### Target stimuli

GRATINGS (contrast, 5 stimuli). These stimuli are horizontal gratings (but not quite sinusoids), with a similar spatial pattern to the middle stimulus in the GRATINGS (density) stimuli from data sets 3 and 4. They were made by convolving horizontal lines spaced every 1.75 deg with a custom band-pass filter. The images were scaled to yield 5 different contrasts of 3%, 10%, 25%, 50% and 100%.

GRATINGS (density; 5 stimuli). These stimuli are similar to horizontal gratings, made by convolving equally spaced horizontal lines with the custom band-pass filter. The 5 stimuli differed in the spacing of the horizontal lines, spaced every 3, 2.5, 1.75, 1, 0.33 deg. The contrast of all stimuli was 25%. The middle stimulus in this sequence was the same as the middle stimulus in the contrast sequence (spacing 1.75 deg, 25% contrast). The highest density (0.33 deg spacing) is close to a sinusoidal grating, as the spacing is the inverse of the peak spatial frequency of the band-pass filter (3 cycles per degree).

SNAKES (contrast, 5 stimuli). The spatial pattern is similar to the snakes stimuli in DS3 and DS4. Contrasts matched the grating contrasts (3%, 10%, 25%, 50% and 100%).

SNAKES (density, 5 stimuli). The spatial pattern is similar to the snakes stimuli in DS3 and DS4, but with 5 different densities of the contours. The contrast for all stimuli was 25% (lower than the contrast of the corresponding stimuli in DS3 and DS4). The range of densities used here was also lower than the range used in DS3 and DS4, with the densest pattern here similar to the middle stimuli in DS3 and DS4.

### Additional stimuli

GRATINGS (orientation, 4 stimuli). These stimuli are the same as the third stimulus in the GRATINGS (density) group (25% contrast, 1.75 deg spacing between contours), except that they are rotated by 0, 45, 90, or 135 deg. Because the 0 deg rotation does not change the image, a new stimulus was not created; for visualization of results, the BOLD measurements and model predictions are plotted for both groups.

GRATINGS (cross, 4 stimuli). These stimuli contain horizontal contours similar to two of the stimuli in the GRATINGS (density) sequence, except that they have periodic vertical blank regions which interrupt the contours. For two of the stimuli, the spacing of the horizontal contours matches the densest stimuli in the density sequence (spacing of 0.33 deg) and for two of the stimuli, the spacing matched the middle stimulus in the density sequence (1.75 deg spacing). In all 4 images, the horizontal contours are interrupted by vertical blanks spaced every 1.75 deg. The vertical blanks are either thick (50% duty cycle; 1st and 3rd stimulus) or thin (25% duty cycle; 2nd and 4th stimuli).

NOISE BARS (density, 5 stimuli). These stimuli have the same contrast apertures as the GRATINGS (density) stimuli. Specifically, there are horizontal bands containing contrast patterns, spaced the same as the grating stimuli (bands every 3, 2.5, 1.75, 1, or 0.33 deg). These stimuli differ from the gratings in that each band contains band-pass filtered noise, equal in power across orientations, rather than horizontal contours.

NOISE BARS (contrast, 5 stimuli). These stimuli are matched in spatial pattern to the middle density of the NOISE BARS (density) stimuli (horizontal lines, spacing 1.75 deg), but scaled in contrast similar to the grating stimuli (3%, 10%, 25%, 50%, 100%).

NOISE BARS (orientation, 4 stimuli). The orientation sequence rotated the middle stimulus of the NOISE BARS (density) group (spacing 1.75 deg, contrast 25%) by 0, 45, 90, or 135 deg.

WAVES (density, 6 stimuli). These are identical to the snakes (density) stimuli, except that they have been filtered by orientation, such that they only contain power at or near the horizontal.

WAVES (contrast, 5 stimuli). These are identical to the snakes (contrast) stimuli, except that they have been filtered by orientation, such that they only contain power at or near the horizontal.

WAVES (orientation, 4 stimuli). These are identical to the densest stimulus in the snakes (density) group, except that they have been filtered by orientation, with filter centered at either 0, 45, 90, or 135 deg.

#### 4.2.3 Stimuli for data set 2

Data set 2 was collected at NYU. The stimuli were nearly identical to those in data set 1, differing only in the following ways. First, the stimuli were 50% larger (18.75 x 18.75 deg and 600 x 600 pixels, rather than 12.5 x 12.5 deg and 400 x 400 pixels). The difference in size did not entail a difference in spatial frequency: The spatial frequency was matched between the two data sets (meaning that the stimuli were re-made with a larger aperture rather than by re-scaling). Second, those stimuli which were oriented were oriented vertically rather than horizontally. This applies to all GRATING stimuli, as well as NOISE BARS and WAVES. Third, the WAVES (density) stimuli had only 4 densities rather than 6. We reduced the number of stimuli to slightly shorten the MRI scans.

### 4.3 MRI

The methods for MRI acquisition and preprocessing for data sets 3 and 4 are described in Kay *et al*. (<u>2013</u>). In brief, each data set comes from one subject, who viewed a variety of stimuli in an event-related fMRI design. Data set 3 was collected over two scan sessions and each stimulus was presented 6 times. Data set 4 was collected over one scan session and each stimulus was presented 3 times. (Note that both in this paper and the Kay website (http://kendrickkay.net/socmodel/), these two datasets are referred to as data sets 3 and 4. However, the Kay website refers to the stimuli for these two datasets as stimulus set 2 and the subjects themselves as “subject B” and “subject C”. We do not adopt these latter two conventions.)

After preprocessing the data (slice-time correction, co-registration, spatial unwarping), a general linear model was applied using the GLMdenoise toolbox (Kay et al., 2013a). The output of this algorithm includes a coefficient (beta weight) for each stimulus for each voxel solved from the whole fMRI session, as well as 30 bootstrapped estimates of each beta weight (bootstrapping across fMRI runs). The publicly available data (http://kendrickkay.net/socmodel/) are already pre-processed, denoised, and organized by ROI. Specifically, the data we used are in the files called “data set03.mat” and “data set04.mat” (http://kendrickkay.net/socmodel/dataset03.mat, http://kendrickkay.net/socmodel/dataset04.mat). The datasets are described on the website as “Dataset 3 (subject B)” and “Dataset 4 (subject C),” respectively. Within the MATLAB files, we used the stored 3D array called “betas” (voxels x stimuli x bootstraps), limited to V1, V2, V3 as indicated in the grouping variables “roi” and “roilabels”, and limited to the 39 stimuli indicated in Supplementary Table 2. Visual areas were identified by retinotopic mapping in a separate session.

#### 4.3.1 Acquisition of data sets 1 and 2

Datasets 1 and 2 were acquired in one scanning session each. Each scanning session had 12 fMRI runs of 249 s each (data set 1) or 241.5 s each (data set 2). For each data set, half of the stimuli were assigned to odd fMRI runs and half to even runs, so that each stimulus was shown 6 times in the session. The stimulus events were 3 s long, consisting of 9 alternations between stimulus exemplar and blank, ⅙ s each. Trial onsets were every 7.5 s (so 4.5 s blank between trials). To help estimate the hemodynamic response function, there were 12 s of blank at the beginning and end of each run, as well as 5 additional trials randomly interspersed with no stimulus (meaning that 5 times during the scan, trials were separated by 15 s instead of 7.5 s). Thus, each complete run consisted of either (25 stimuli + 5 blanks) * 7.5 s + 24 s = 249 s (data set 1) or (24 stimuli + 5 blanks) * 7.5 s + 24 s = 241.5 s (data set 2).

All MRI data were acquired at New York University Center for Brain Imaging using a Siemens Allegra 3T head-only scanner with a Nova Medical phased array, 8-channel receive surface coil (NMSC072). For each participant, we collected functional images (single shot echo planar images, 1500 ms TR, 30 ms TE, and 72° flip angle). Voxels were 2.0 mm^3^ isotropic, with 24 slices, with an inplane sampling of 104 x 80 voxels (208 mm A/P x 160 mm L/R). The slice prescription covered most of the occipital lobe, and the posterior part of both the temporal and parietal lobes. Images were corrected for B0 field inhomogeneity using a calibration scan and Center for Brain Imaging algorithms during offline image reconstruction.

We also acquired 1 or 2 T1-weighted whole-brain anatomical scans (MPRAGE sequence; 1mm^3^), as well as a T1-weighted “inplane” image with the same slice prescription as the functional scans. This scan had an inplane resolution of 1.25 X 1.25 mm and a slice thickness of 2.5 mm, and was collected to aid alignment of the functional images to the high-resolution T1 weighted anatomical images.

In a separate session, retinotopy scans were collected and analyzed using a pRF model as implemented in the Vistasoft software tool (https://github.com/vistalab/vistasoft). The methods for acquisition and analysis of the retinotopy data are identical to that described by Zhou et al. (2018).

#### 4.3.2 Data Preprocessing and Analysis

Data preprocessing. Processing of the fMRI data was identical to that described by Zhou et al. (2018): We coregistered and segmented the T1 weighted whole-brain anatomical images into gray and white matter voxels using FreeSurfer’s autosegmentation algorithm (http://surfer.nmr.mgh.harvard.edu). Using custom software Vistasoft (https://github.com/vistalab/vistasoft), the functional data were slice-time corrected by resampling the time series in each slice to the center of each 1.5 s volume. Data were then motion-corrected by coregistering all volumes of all scans to the first volume of the first scan. The first 8 volumes (12 s) of each scan were discarded for analysis to allow longitudinal magnetization and stabilized hemodynamic response.

*GLM.* The preprocessed fMRI data were then fit by a general linear model, GLMDenoise (Kay et al., 2013a). This algorithm denoises the data by projecting out nuisance regressors derived in a data-driven manner, and estimates coefficients for each of the 48 or 50 stimuli for each voxel in the functional images. The algorithm bootstraps the data over fMRI runs. For data sets 1 and 2, we generated 50 bootstraps for data set 1 and 100 bootstraps for data set 2. The publicly available data from Kay *et al*. (<u>2013</u>), included 30 bootstraps per subject. The algorithm also estimated a hemodynamic impulse response function as a finite impulse response function, with 35 time points (52.5 s) per subject.

*ROIs.* Regions of interest for V1, V2, V3 were delineated manually using the Vistasoft (https://github.com/vistalab/vistasoft) graphical user interface to visualize the results of the pRF models. These methods for identifying these boundaries are well established, as described in many publications (Engel et al., 1997; Sereno et al., 1995; summarized by Wandell and Winawer, 2011). The ROIs for V1, V2 and V3 were identified on the cortical surface and then projected to the functional images. For purposes of data summary and model fitting, we took the average signal from each ROI. We did this by averaging the beta weight across voxels within an ROI separately for each stimulus, after voxel selection (**Supplementary Table 1**). Because noise can be correlated across voxels, but should not be correlated across scans, when we bootstrapped the data, we average across voxels within an ROI for each bootstrap. For the purposes of model fitting, each of the 4 datasets comprised two matrices, one for the means and one for the standard deviation across bootstraps, each of which had a size equal to the number of stimuli by number of ROIs.

### 4.4 Model Equations

In the Results, we compared the accuracy of four models fit to the data, three of which are based on existing models or empirical findings–a contrast energy model, a untuned normalization model, and an orientation tuned normalization model–and one new model, which computes normalization by orientation anisotropy. In this section we describe the computation that comprises each model.

All four models consist of three primary steps: (1) computation of oriented contrast energy, (2) pooling across orientation and space, and (3) a power-law nonlinearity. Steps 1 and 3 are identical for all models. Step 2, spatial pooling, varies between models.

*1. Contrast energy:* We denote by 𝐼(𝑥, 𝑦) the value of the pre-processed input image at coordinates (𝑥, 𝑦). The pre-processing causes the image values to have mean 0 and range from -0.5 to 0.5. The image is projected onto a set of 64 Gabor filters, which comprise 8 orientations 𝜃, 4 spatial frequencies 𝑓, and 2 phases 𝜙, separated by 90° (i.e., “quadrature”). The orientations are sampled from 0 pi to 7/8 pi with a 1/8 pi interval; the filters’ spatial frequencies are 0.75, 1.5, 3, 6 cycles per degree (cpd).

𝐹(𝑥, 𝑦, 𝜃, 𝑓, 𝜙) indicates the Gabor filter at a spatial location (𝑥, 𝑦), orientation 𝜃, spatial frequency 𝑓, and phase 𝜙. Each filter had a total of 6 cycles each. The outputs over the two phases are squared and summed to compute the contrast energy and summed across spatial frequencies. Finally, the contrast energy as a function of spatial position (𝑥, 𝑦) and orientation 𝜃 becomes

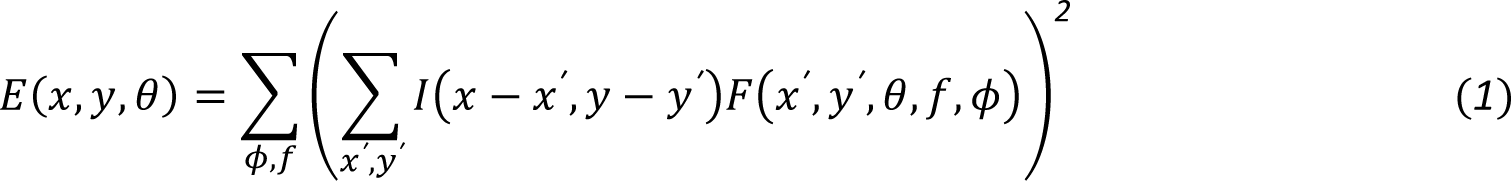

We convolve the image 𝐼 and the filter 𝐹. The computation of contrast energy has no free parameters. Prior to convolution, stimuli were padded with uniform gray (mean luminance) on all sides by the width of the largest filter. After convolution, all energy images were downsampled to 12 pixels per degree for computational efficiency.

*2. Spatial pooling.* Each model differs in how contrast energy is pooled to yield a scalar value, 𝑠:

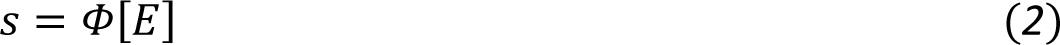

where we use square brackets to indicate a function of a function (also called a *functional*). We describe the pooling functional 𝛷, for each model below. **Error! Reference source not found.Error! Reference source not found.Error! Reference source not found.Error! Reference source not found.Error!** Reference source not found.Error! Reference source not found.Error! Reference source not found.

*3. Power-law nonlinearity.* Finally, the scalar is passed through a power-law nonlinearity to predict the BOLD amplitude 𝑟 in units of percent signal change:

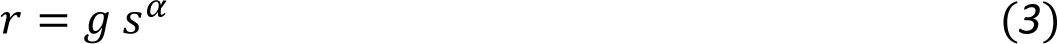

Where 𝑔 is the gain and 𝛼 is the exponent parameter. These are free parameters fit to the fMRI data. The power-law nonlinearity is similar to divisive normalization in the case where each unit in a population is normalized by the same pool (Kay et al., 2013b).

#### 4.4.1 Pooling functional for contrast energy model

In the contrast energy (CE) model, the contrast energy is summed over orientations and space to yield a scalar output, 𝑠.

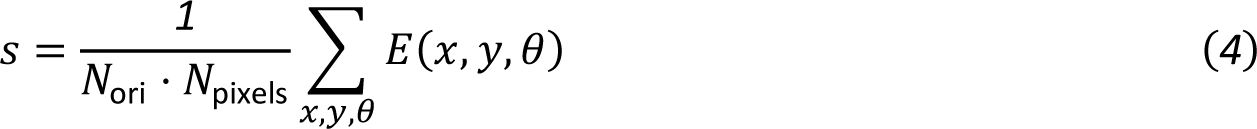

𝑁_ori_ is the number of orientation channels (always 8) and 𝑁_pixels_ is the number of pixels per stimulus in the padded images (344^2^, 419^2^, or 342^2^). There are no free parameters in this pooling functional for contrast energy, so the complete contrast energy model has only two free parameters, 𝑔 and 𝛼, both from the power-law nonlinearity step (**Eq. 3**).

#### 4.4.2 Pooling functional for divisive normalization model

The contrast energy in the untuned divisive normalization (DN) model is normalized before it is summed: Each (𝑥, 𝑦, 𝜃) element in the energy image is normalized by a weighted sum of elements at (𝑥*^′^*, 𝑦*^′^*, 𝜃*^′^*). The weighting is a Gaussian function of distance from location (𝑥, 𝑦) and can thus be expressed as a convolution of the contrast energy 𝐸, with a Gaussian, 𝐺:

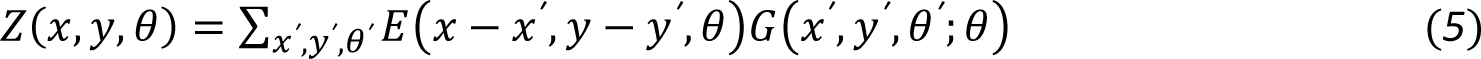

The standard deviation of 𝐺 is 4% of the padded image size, which is approximately 1 deg. 𝐺 is identical across the 8 orientations.

The normalized contrast energy is

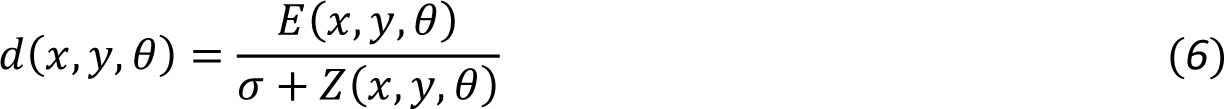

where 𝜎 is a parameter to control the strength of normalization. When 𝜎 is large, the normalization is low, and the overall expression approximates the contrast energy model. When 𝜎 approaches 0, there is strong normalization. We then sum 𝑑 across space and orientation to result in the scalar, 𝑠.

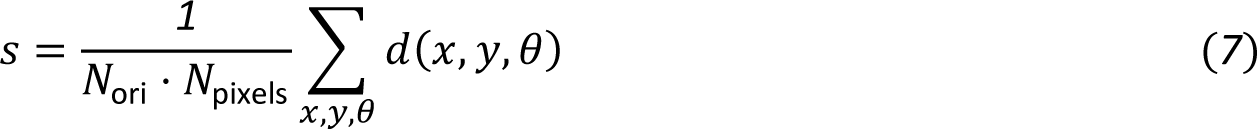

The pooling functional for divisive normalization introduces one free parameter, 𝜎. As with the contrast energy model, a power-law nonlinearity is applied to 𝑠 to predict the BOLD response in percent signal change (**Eq. 3**). Hence the complete model has three free parameters.

We note that the complete divisive normalization model has two similar non-linearities, one in the pooling functional (divisive normalization) and one on the final output (power-law). This is consistent with prior work showing that two stages of normalization (a cascade model) improved model accuracy (<u>Kay, Winawer, Rokem, et al. 2013</u>).

#### 4.4.3 Pooling functional for orientation-tuned normalization model

We implement the orientation-tuned normalization (OTN) model identically to the divisive normalization model (**Eqs 5-7**), except that the contrast energy normalizer 𝑍(𝑥, 𝑦, 𝜃) is now orientation-tuned. When the orientation channel of the image and filter matches, 𝜃′ = 𝜃, the 2-D filter of the channel is a 2D Gaussian identical to the untuned normalization (4.4.2). At all other orientations (i.e., 𝜃′ ≠ 𝜃), the filter is a symmetric 2D Gaussian distribution with a much small standard deviation, effectively just one pixel (**Figure 4**). This is akin to summing two forms of normalization, cross-orientation suppression (same location, other orientations), and an orientation-tuned surround (same orientation, other locations). As with the untuned normalization model, the pooling functional introduces only one free parameter, 𝜎. The complete model, including the power-law non-linearity (**Eq. 3**) has 3 free parameters.

#### 4.4.4 Pooling functional for normalization by orientation anisotropy

In the normalization by orientation anisotropy (NOA) model, the pooling step first sums the contrast energy across space within an orientation band, resulting in one value per orientation band, 𝐸_ori_(𝜃):

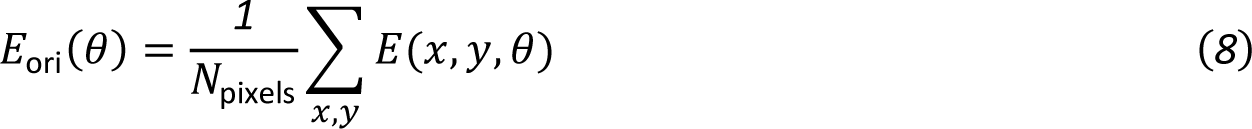

𝐸_ori_ indicates the oriented energy. This energy at each orientation is then normalized by the standard deviation across the 8 orientations, and then summed to produce a scalar.

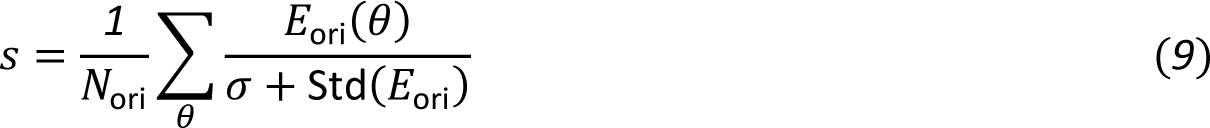

Where 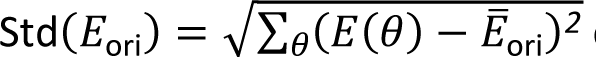 calculates the standard deviation of the oriented energy and a non-negative parameter 𝑤 controls the strength of the normalization. When 𝜎 is large, the normalization is low, and the overall expression approximates the contrast energy model. When 𝜎 approaches 0, there is strong normalization by the standard deviation across orientation channel outputs. Calculating the standard deviation of oriented energy involves a squaring operation. To keep the parameters comparable across different models, we also square the numerator and the parameter 𝜎. The NOA pooling functional has one free parameter, 𝜎. The complete model, including the power-law non-linearity (**Eq. 3**) has 3 free parameters.

### 4.5 Optimization

In each model, we fitted the model free parameters using the MATLAB optimization tool *fmincon* by minimizing the squared error between the model prediction and the corresponding BOLD amplitude. Because each stimulus consisted of 9 exemplars shown to the subject in rapid succession, the model prediction for each stimulus was obtained by averaging the model predictions across the exemplars. To avoid getting stuck in the local minima of the nonconvex landscape, we ran the optimization algorithm with 40 different parameter initializations. Each initialized value was picked randomly. All parameters were unbounded in the search, minimizing human interference in the fitting. Parameter 𝛼 is passed through a sigmoid function to ensure its value is between 0 and 1.

### 4.6 Cross-validation scheme

All models were fit using an *n*-fold leave-one-out cross-validation scheme, where *n* is the number of stimuli. Thus, the BOLD signal prediction for each stimulus was generated by a model fit to all stimuli except that one. Under this scheme, the models are less likely to overfit data sets.

### 4.7 Accuracy metric

The model accuracy was quantified as the percentage of the explained variance (𝑅^2^) in the human BOLD data by the cross-validated model predictions,

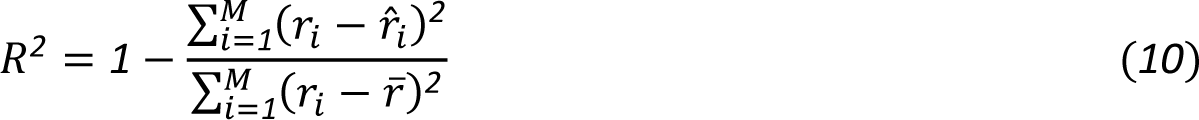

where 𝑟_𝑖_ represents the BOLD amplitude to the *i^th^* stimulus, 𝑟_𝑖_ represents the corresponding model prediction, and r̄ is the mean response across stimuli. We can understand this metric as the extra uncertainty reduction brought by the model beyond describing the BOLD data by its mean.

## Supplementary figures

**Figure S1:**
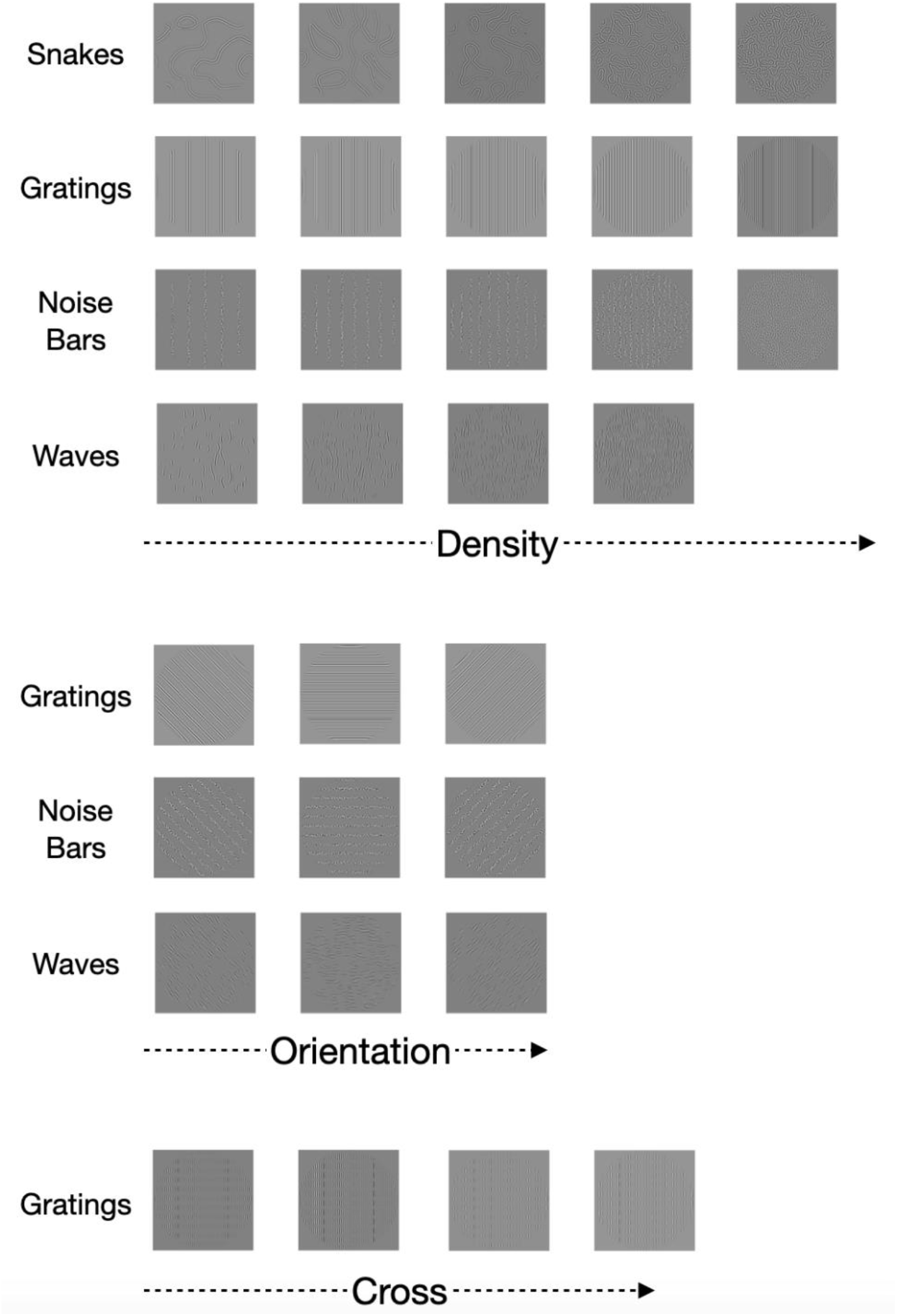
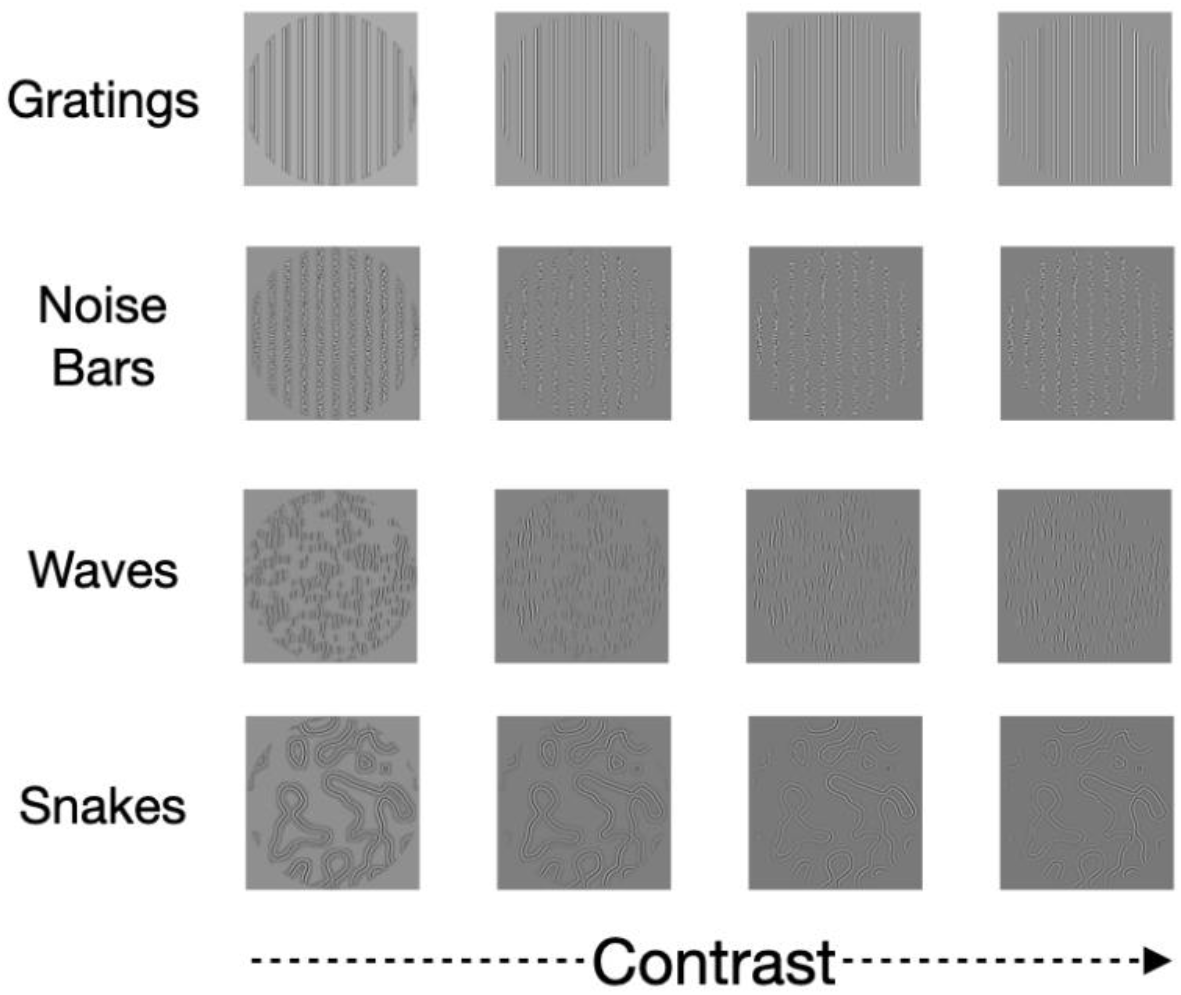
Stimuli for data set 2

**Figure S2a:**
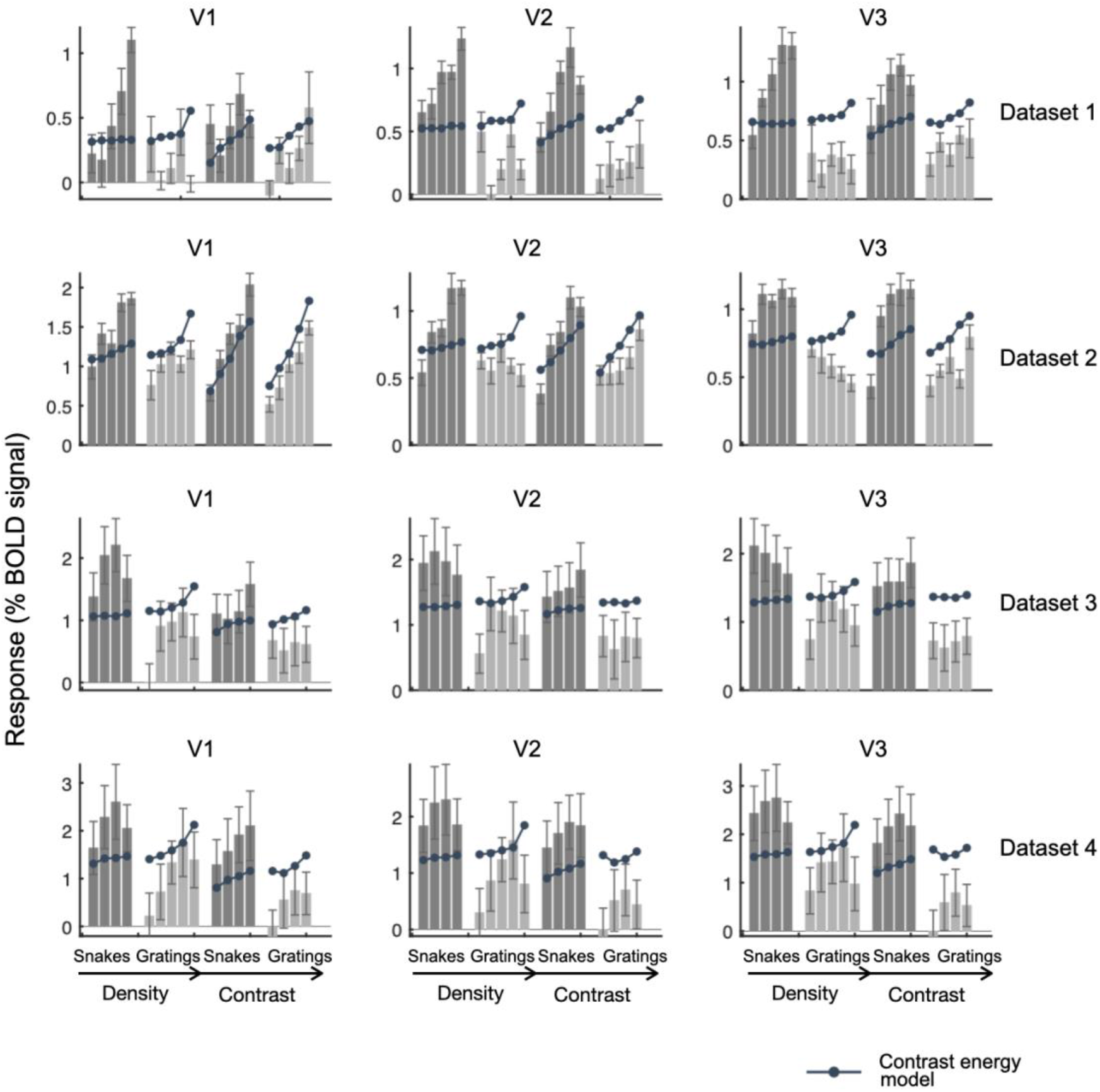
Responses and contrast energy model fits for target stimuli, all datasets

**Figure S2b:**
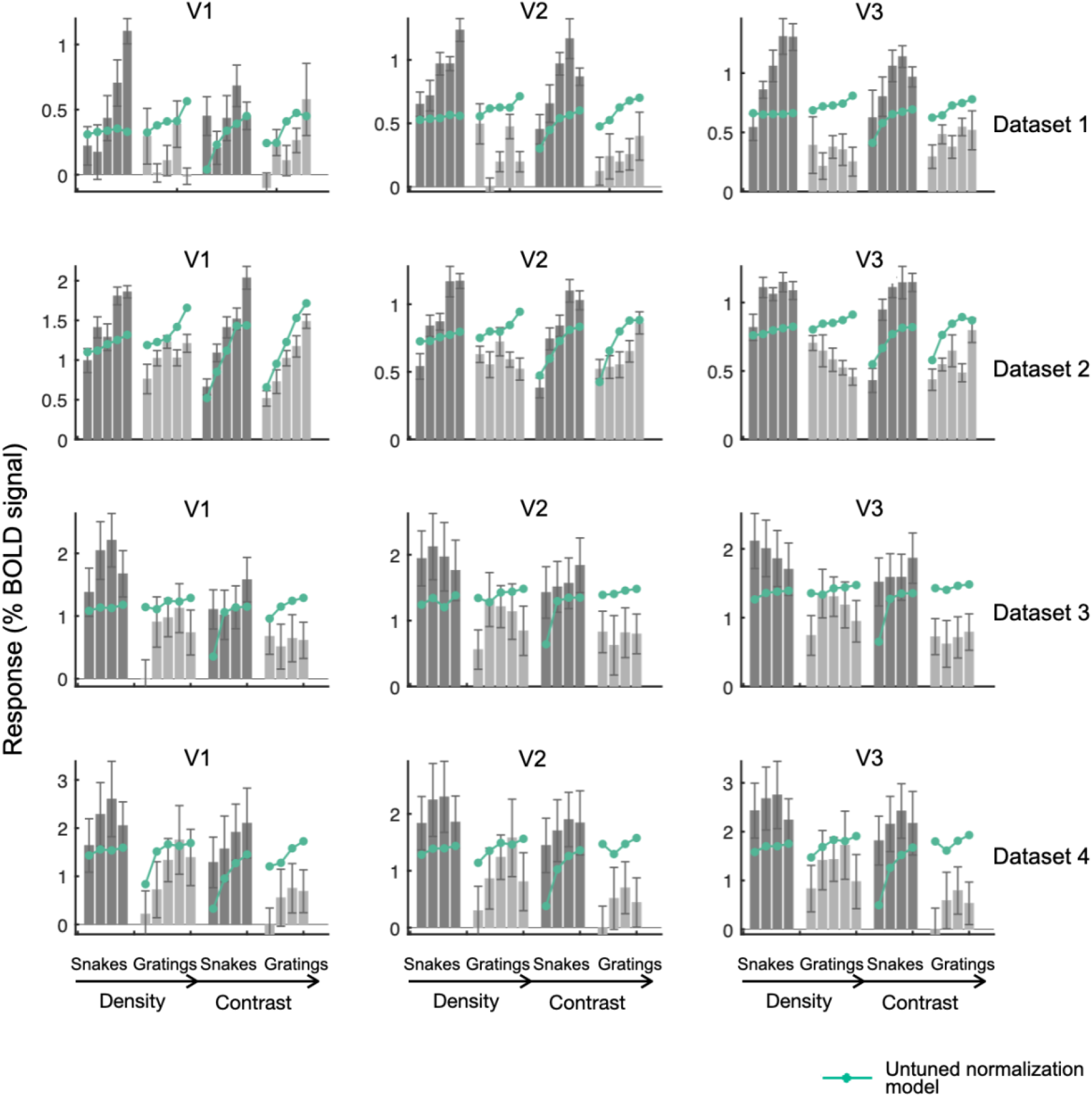
Responses and Untuned normalization fits for target stimuli, all datasets

**Figure S2c:**
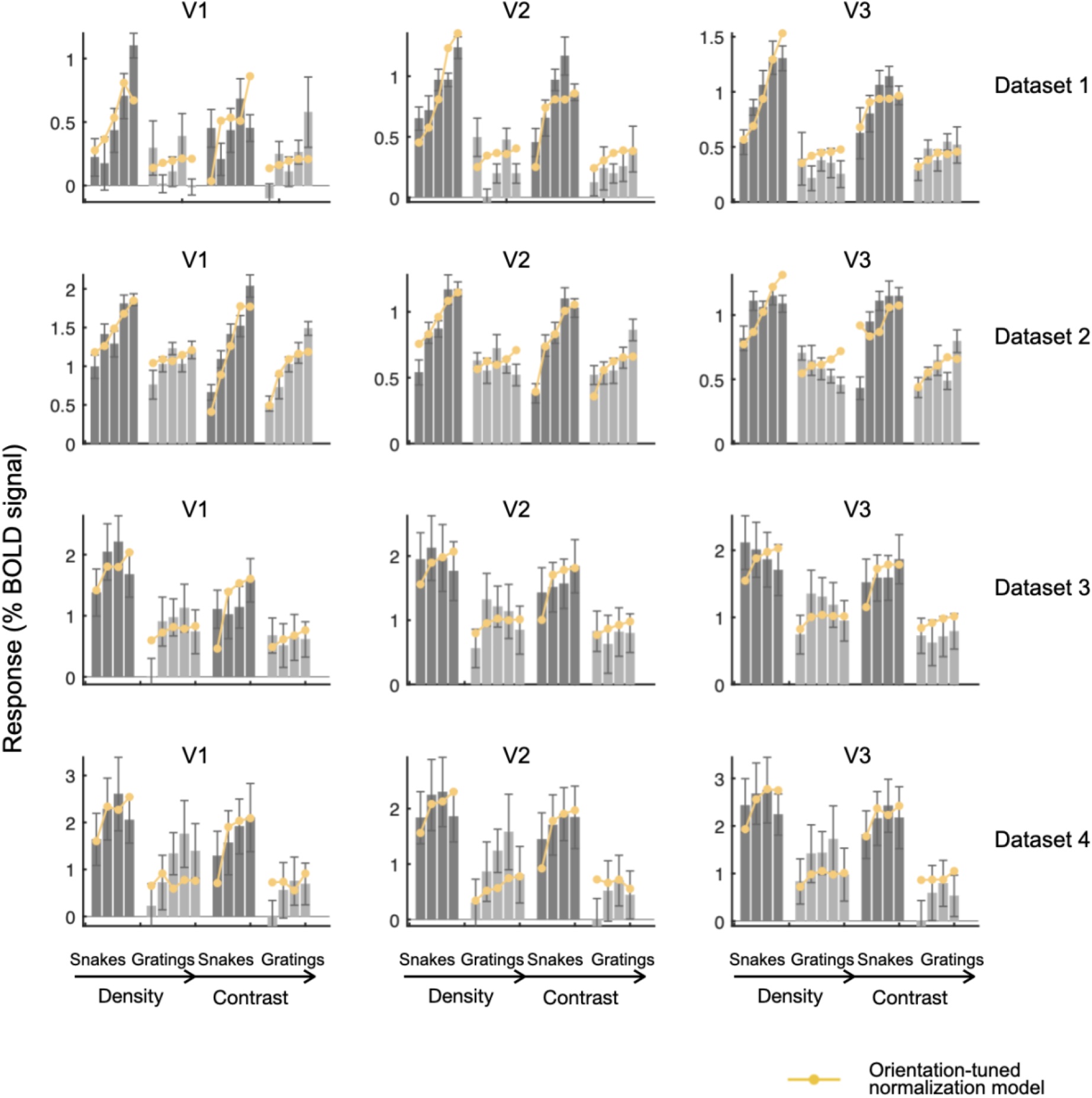
Responses and orientation-tuned normalization model fits for target stimuli, all datasets

**Figure S2d:**
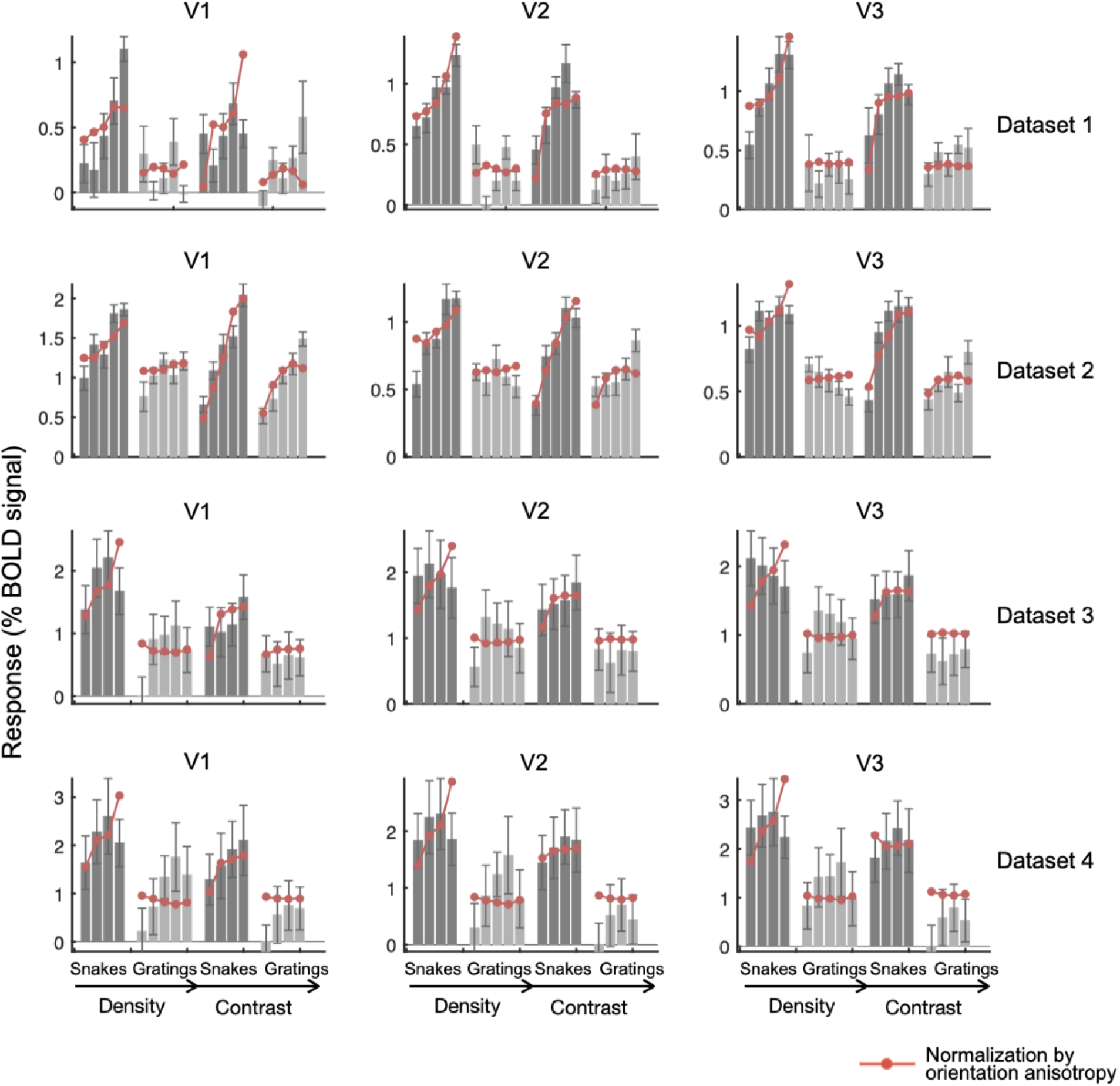
Responses and normalization by orientation anisotropy model fits for target stimuli, all datasets

**Figure S3a:**
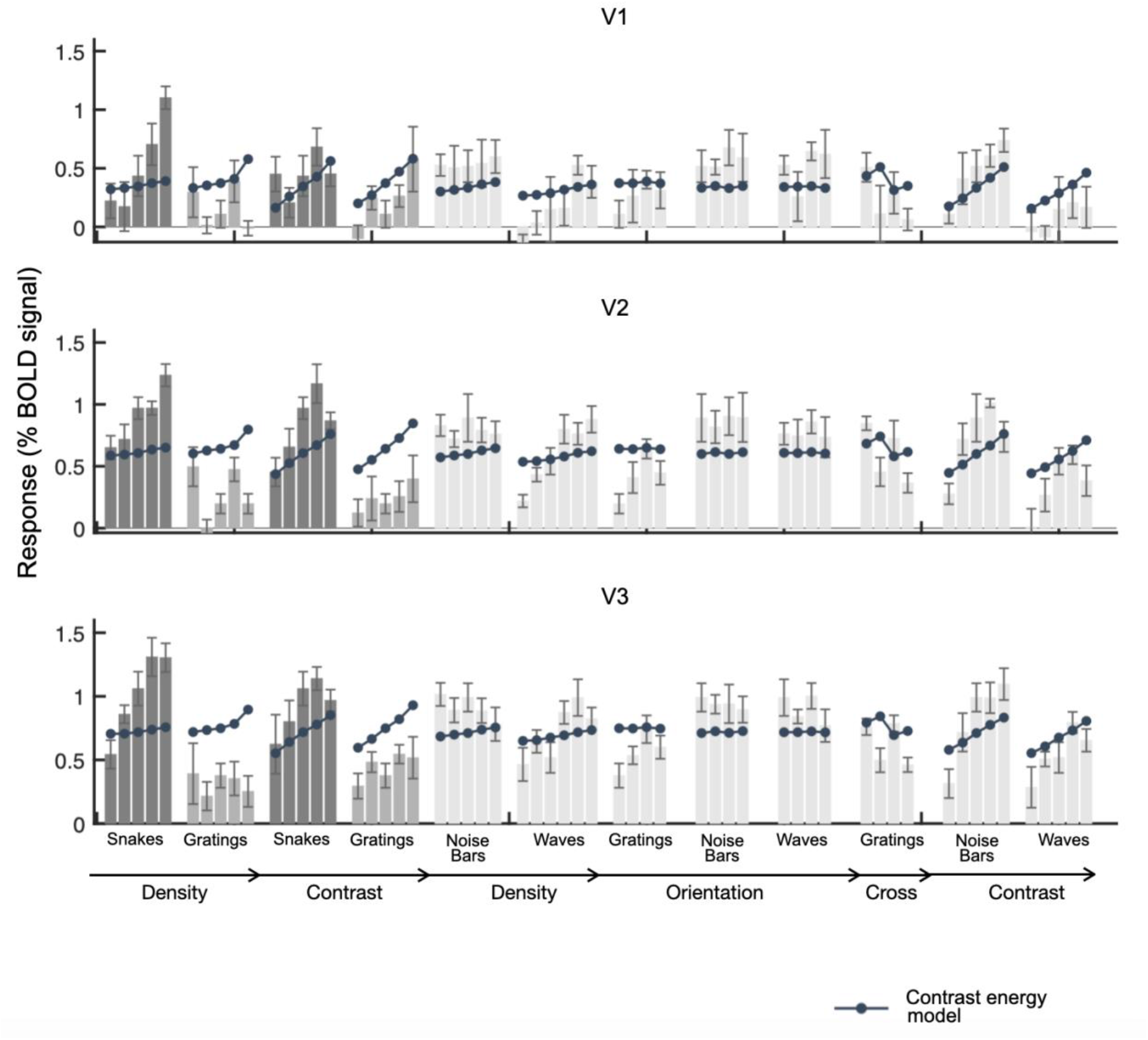
Responses and contrast energy model fits for all stimuli, data set 1

**Figure S3b:**
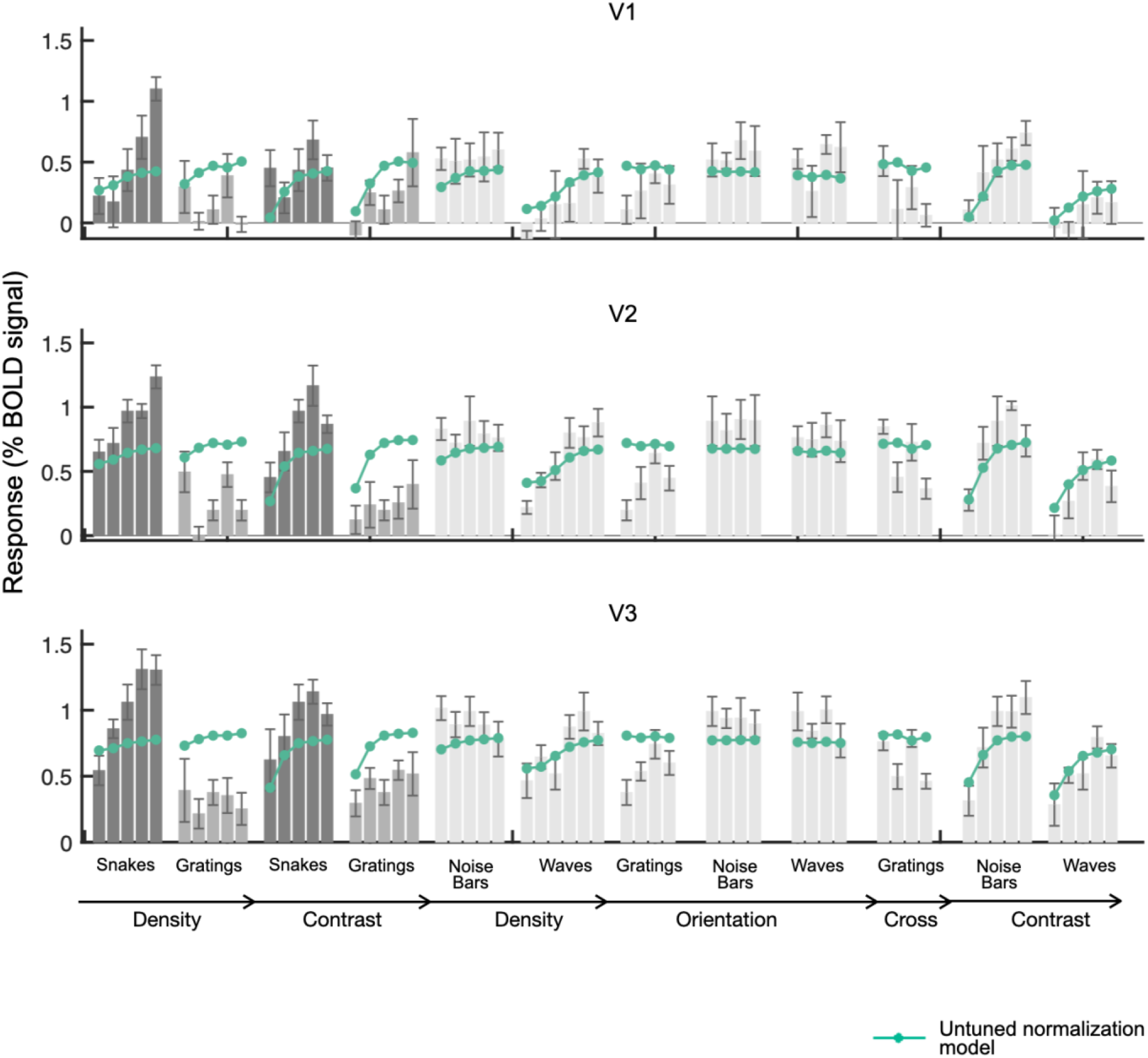
Responses and untuned normalization model fits for all stimuli, data set 1

**Figure S3c:**
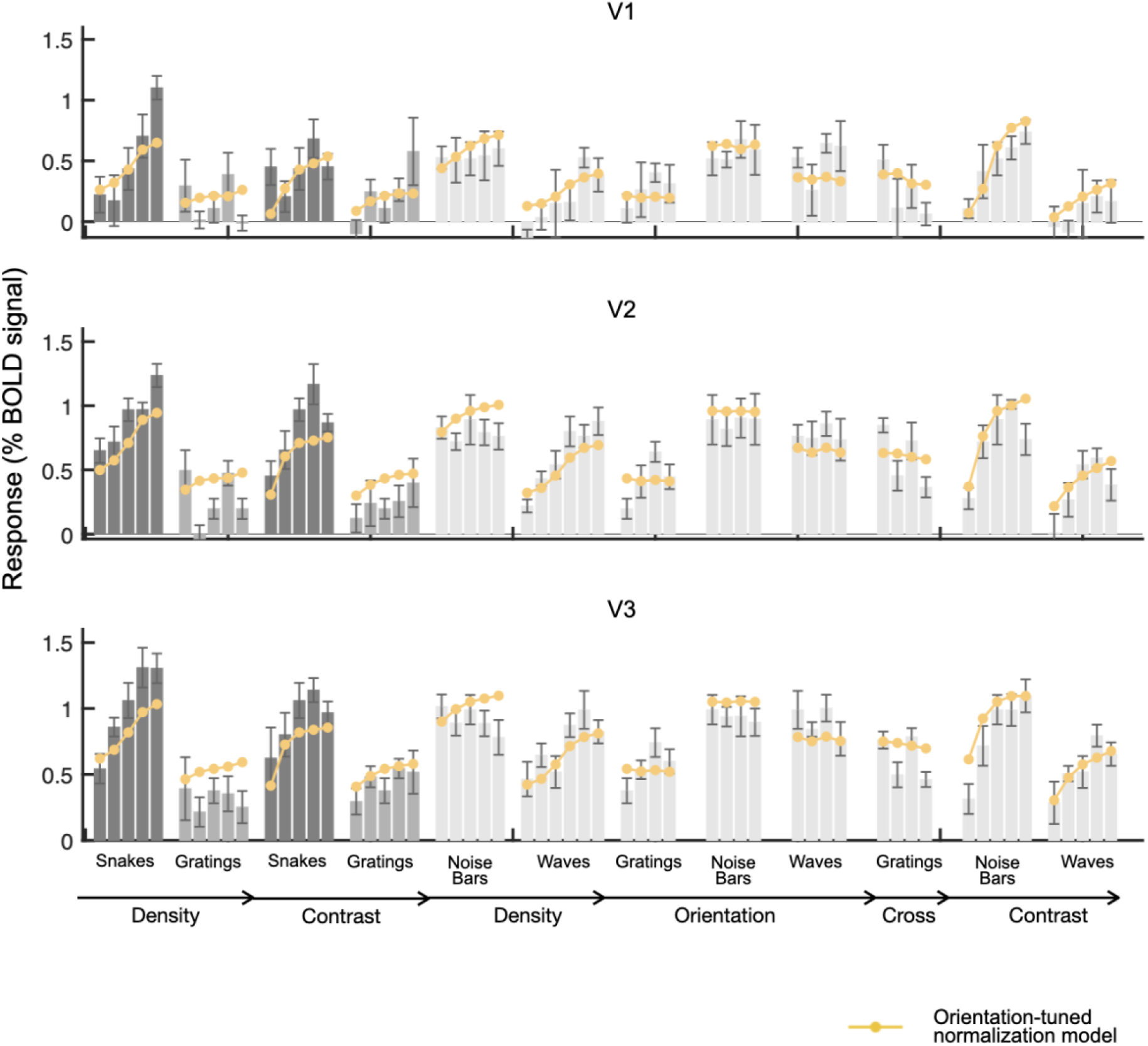
Responses and orientation-tuned normalization model fits for all stimuli, data set 1

**Figure S3d:**
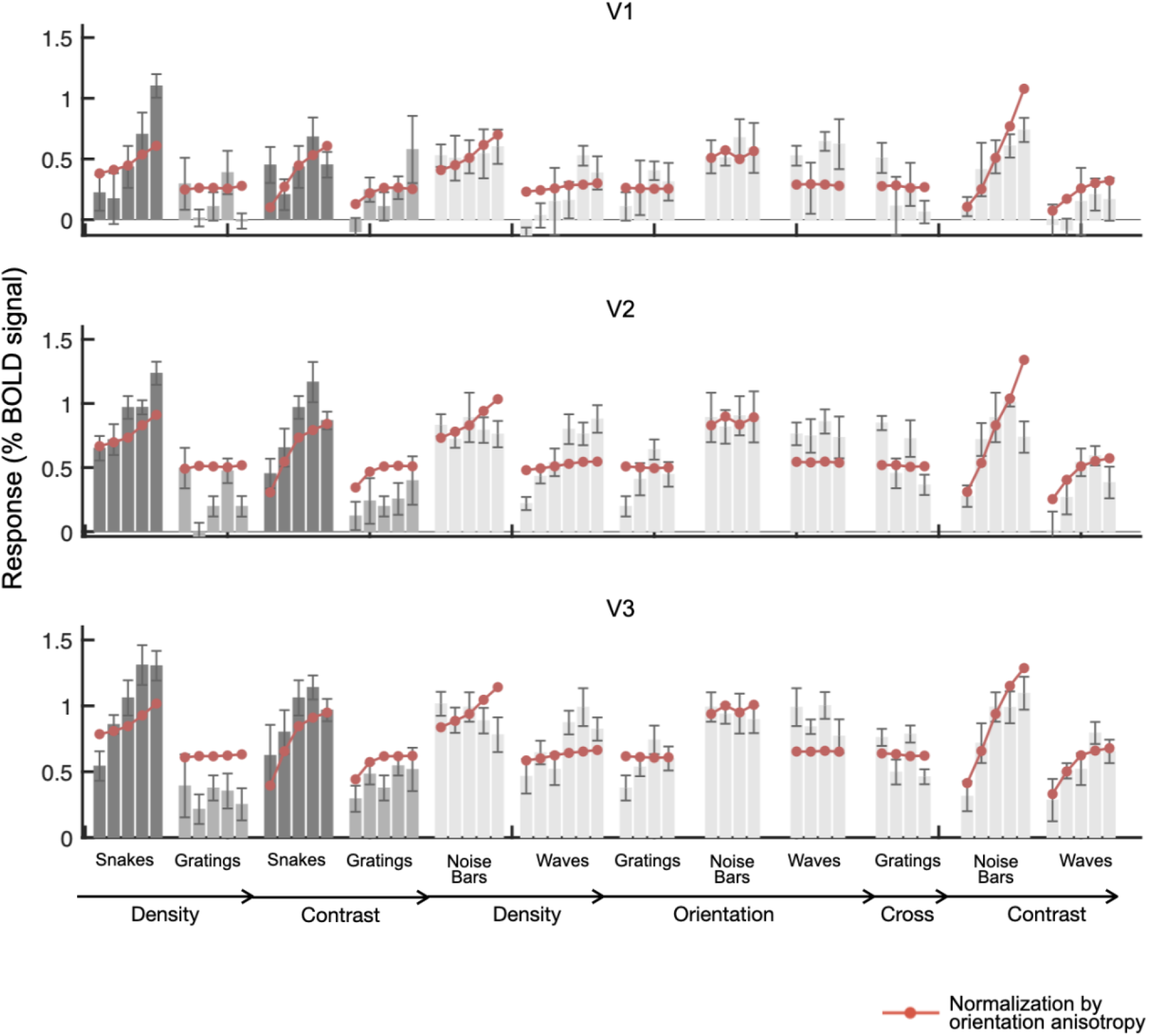
Responses and normalization by orientation anisotropy model fits for all stimuli, data set 1

**Figure S4a:**
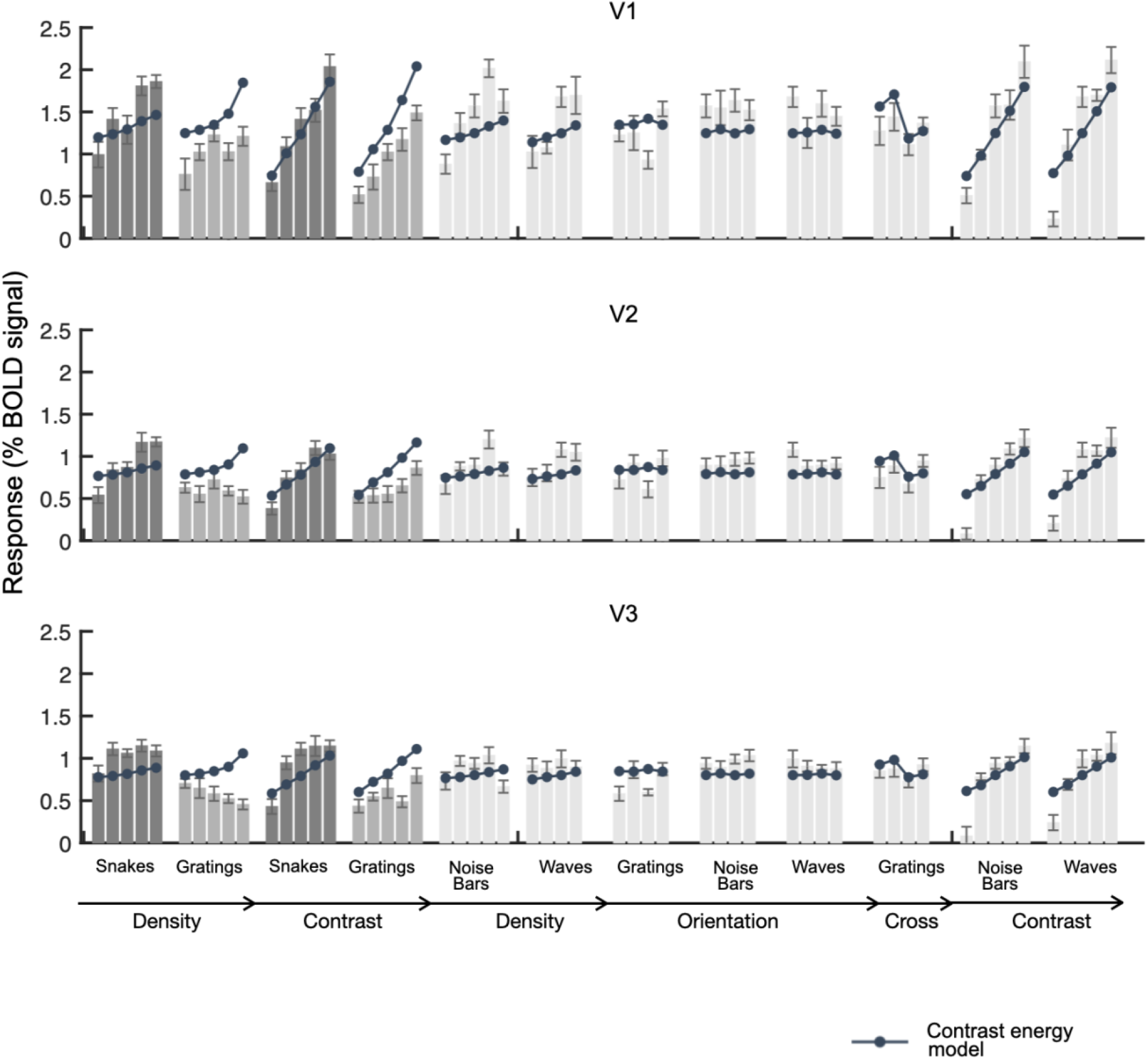
Responses and contrast energy model fits for all stimuli, data set 2

**Figure S4b:**
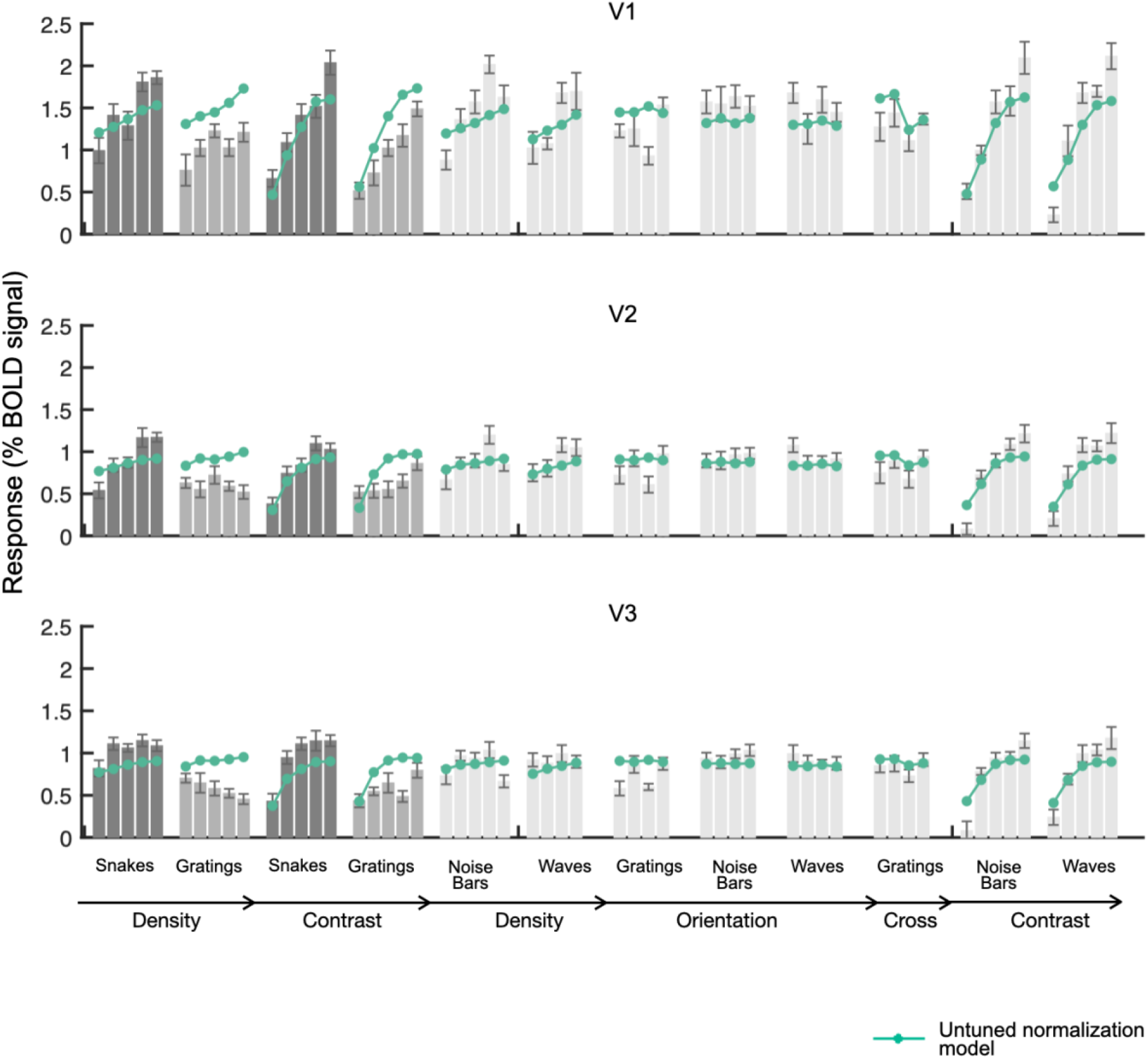
Responses and untuned normalization model fits for all stimuli, data set 2

**Figure S4c:**
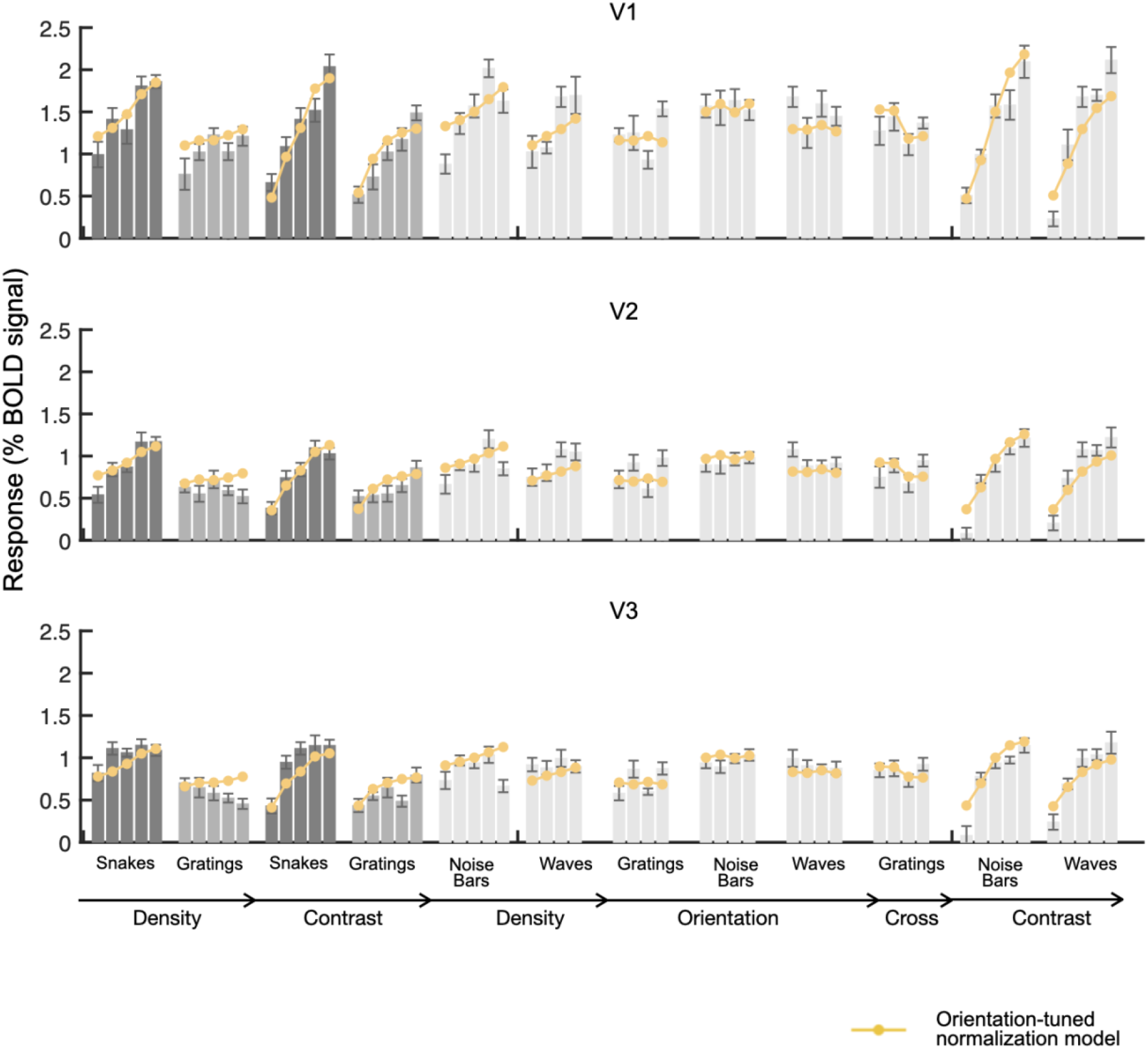
Responses and orientation-tuned normalization model fits for all stimuli, data set 2

**Figure S4d:**
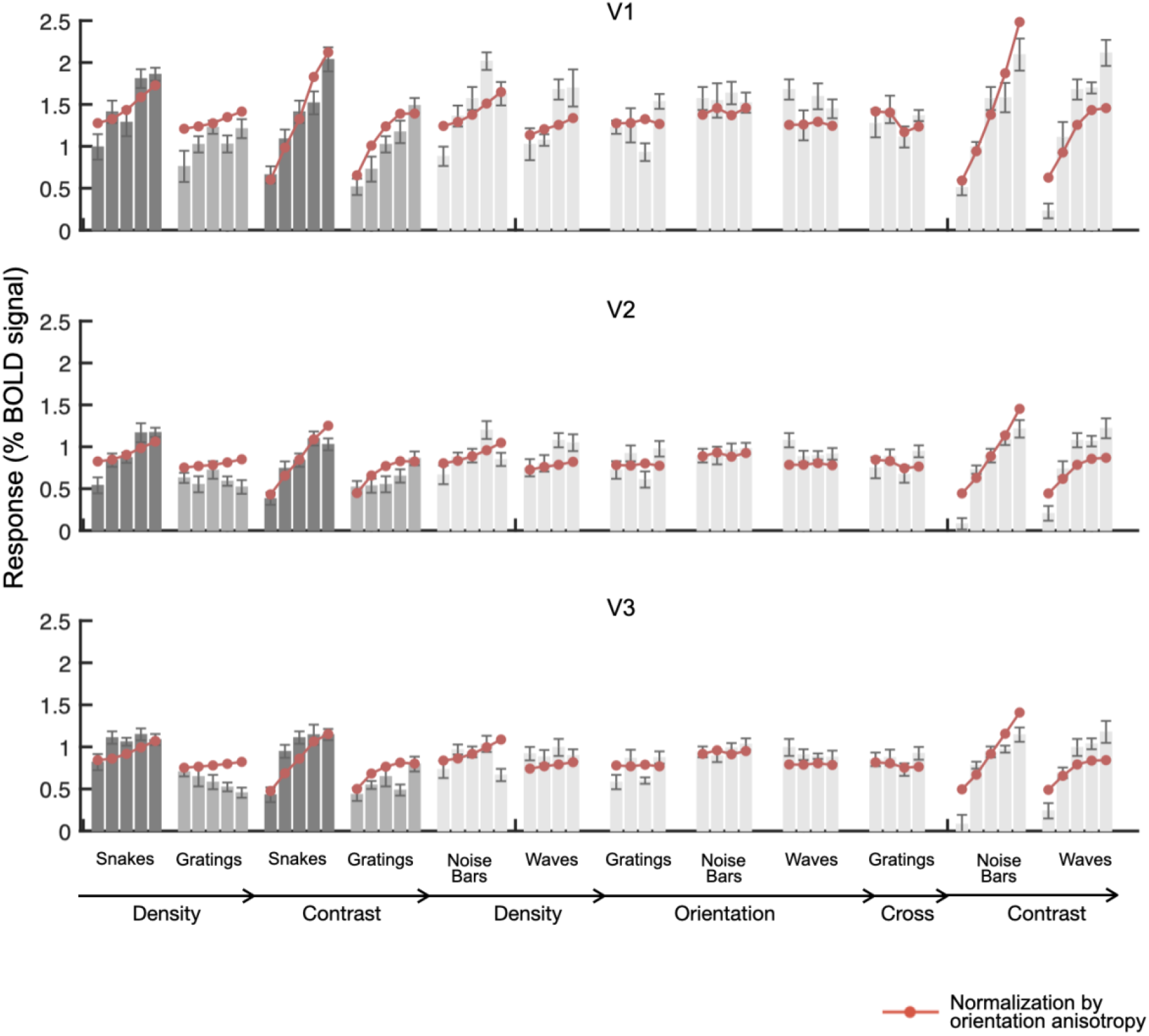
Responses and normalization by orientation anisotropy model fits for all stimuli, data set 2

**Figure S5a:**
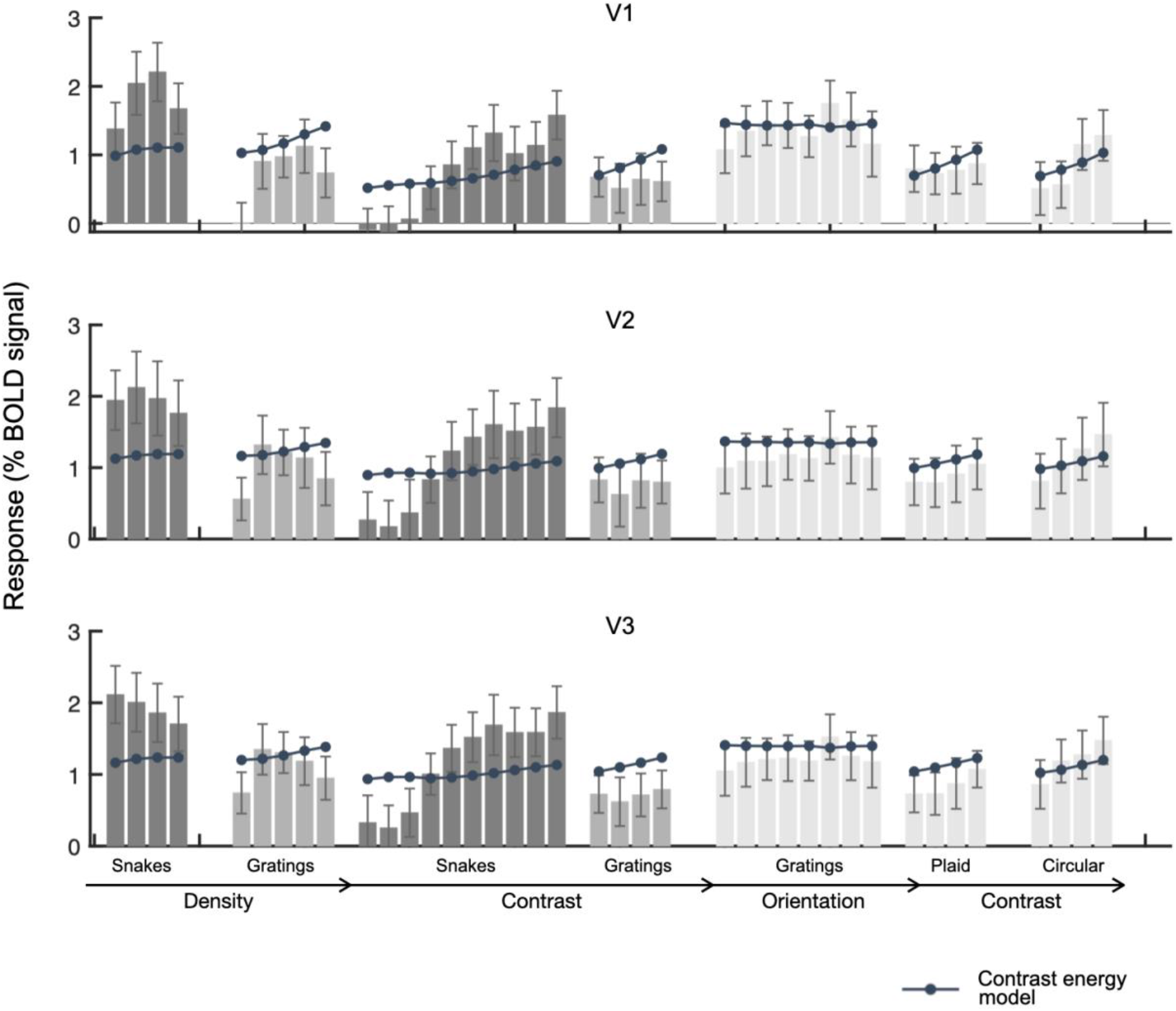
Responses and contrast energy model fits for all stimuli, data set 3

**Figure S5b:**
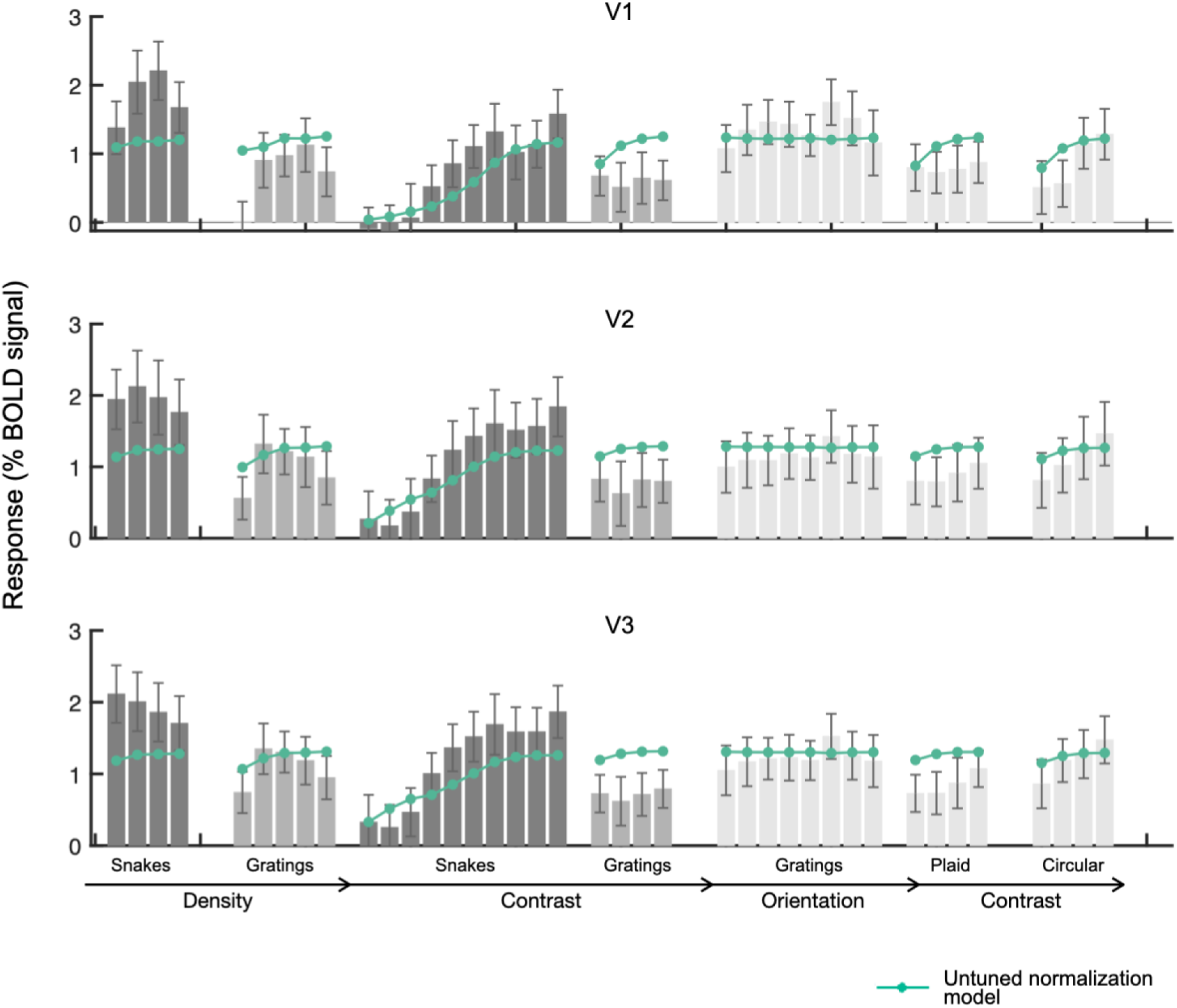
Responses and untuned normalization model fits for all stimuli, data set 3

**Figure S5c:**
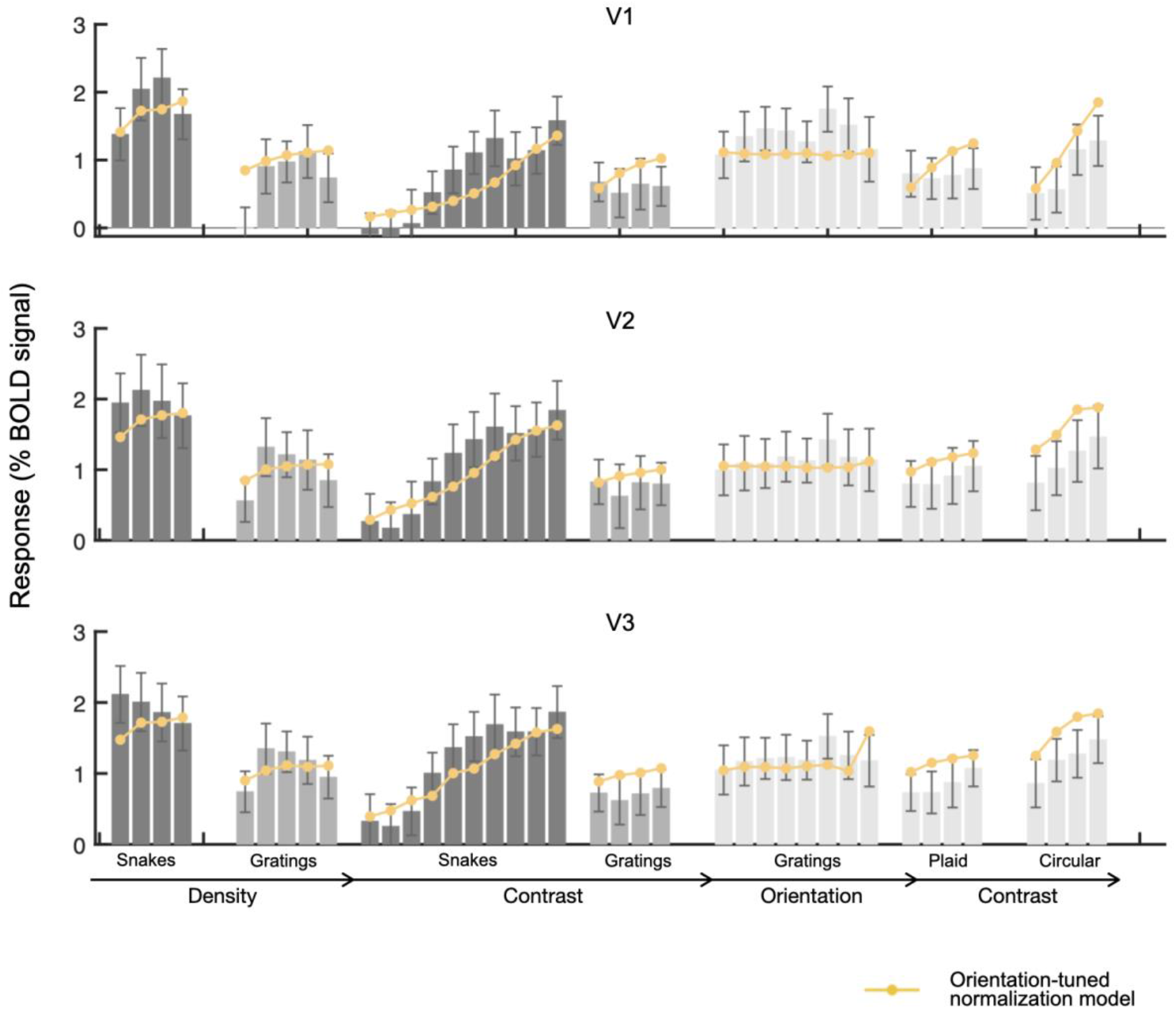
Responses and orientation-tuned normalization model fits for all stimuli, data set 3

**Figure S5d:**
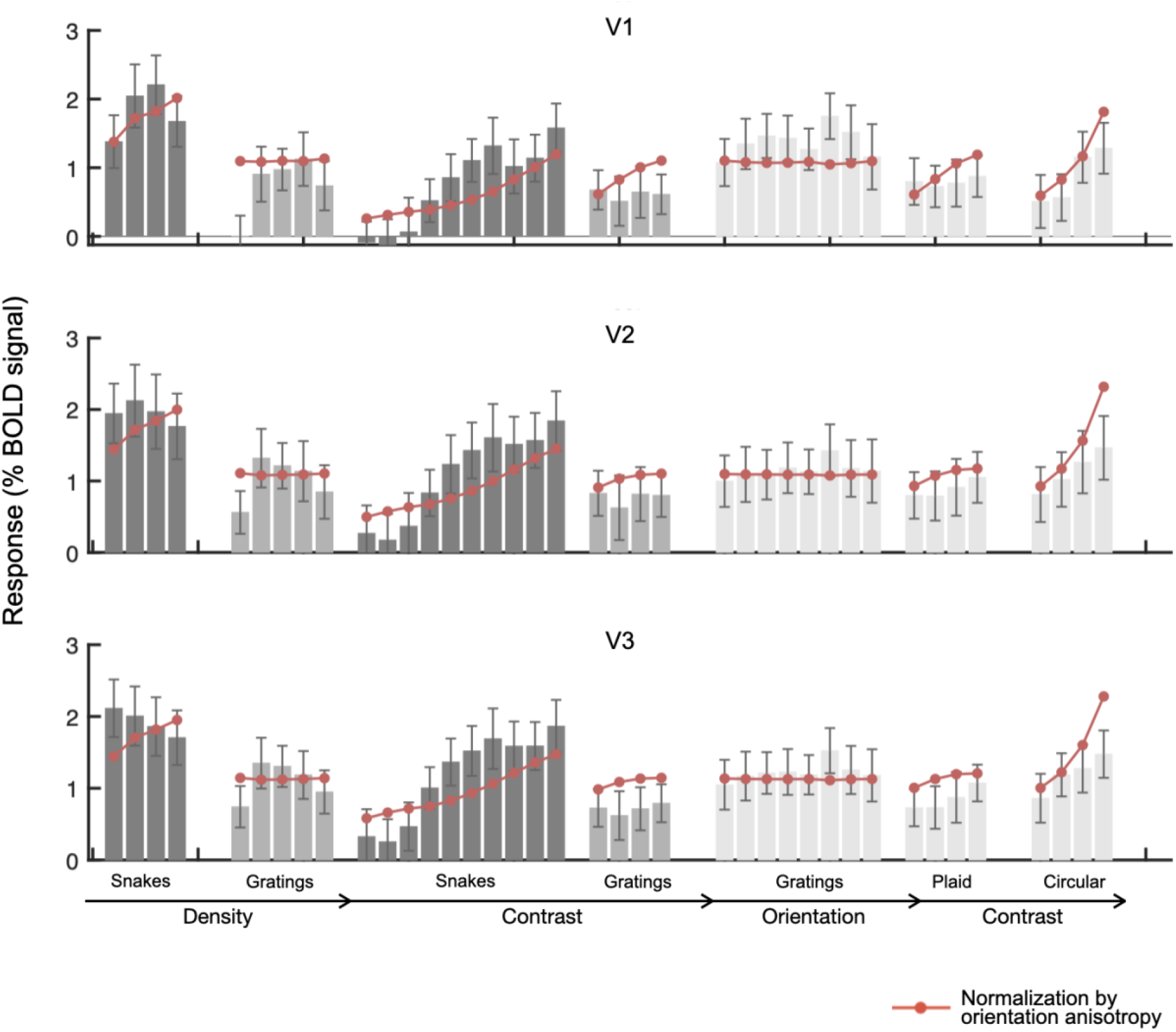
Responses and normalization by orientation anisotropy model fits for all stimuli, data set 3

**Figure S6a:**
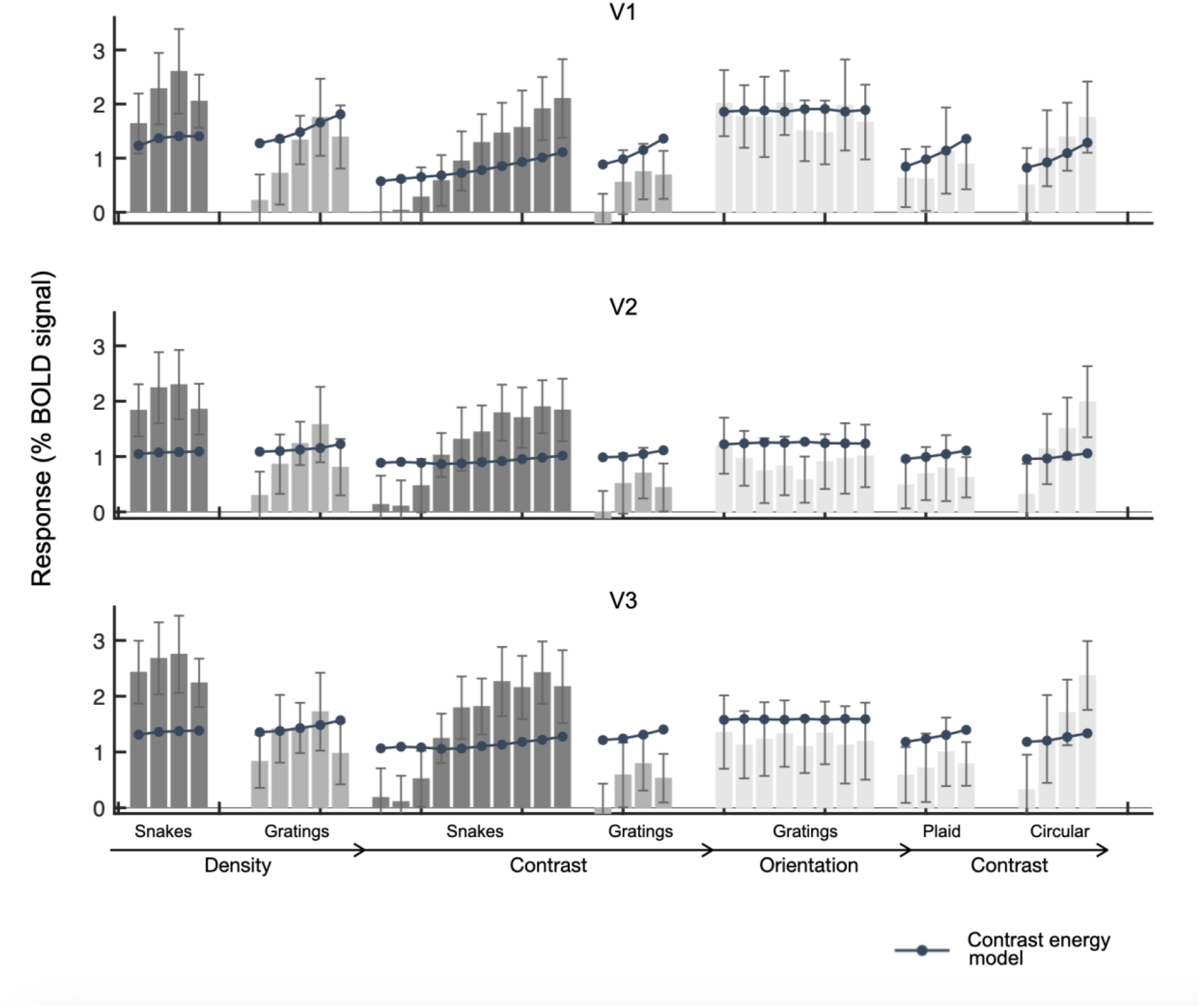
Responses and contrast energy model fits for all stimuli, data set 4

**Figure S6b:**
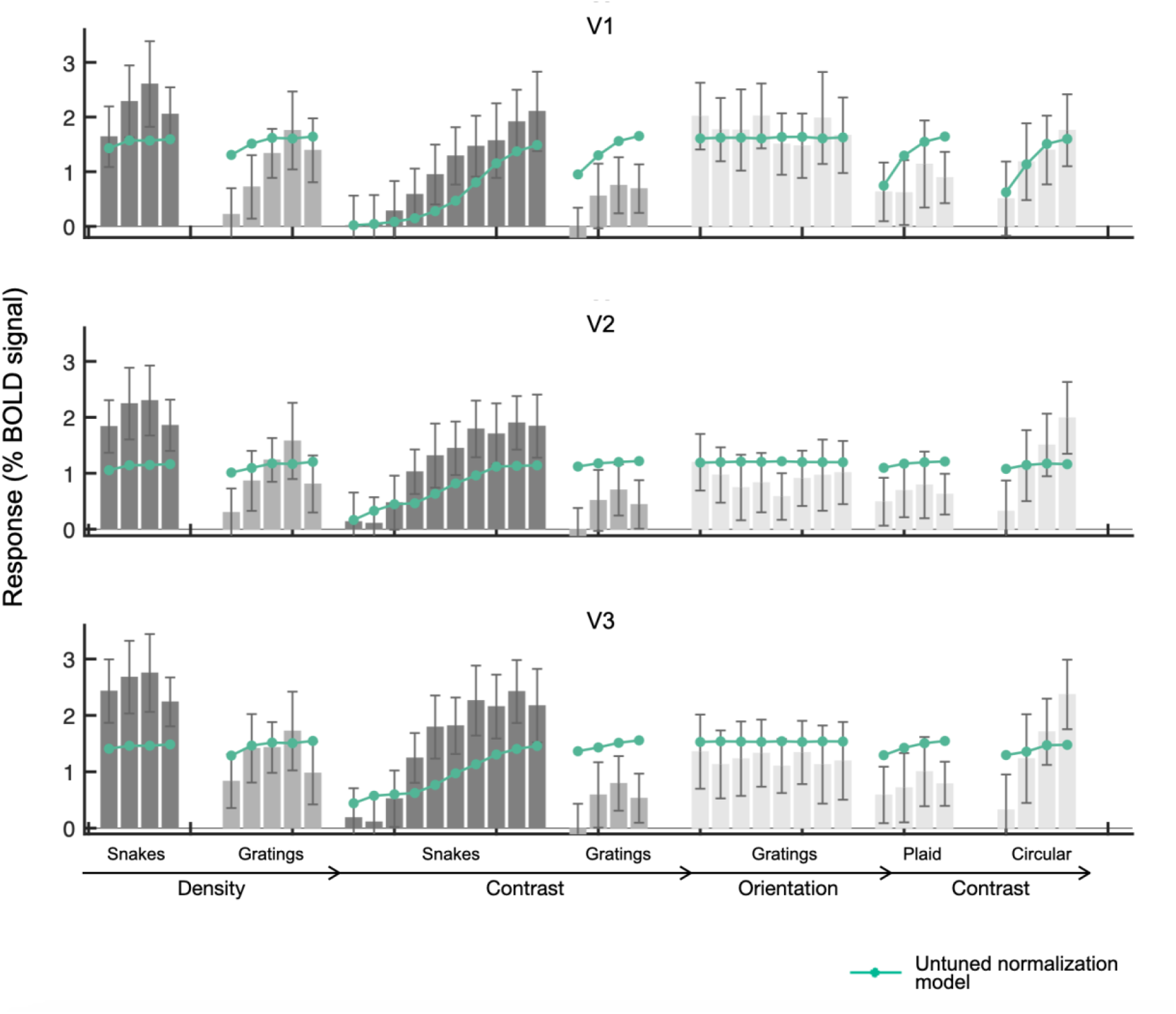
Responses and untuned normalization model fits for all stimuli, data set 4

**Figure S6c:**
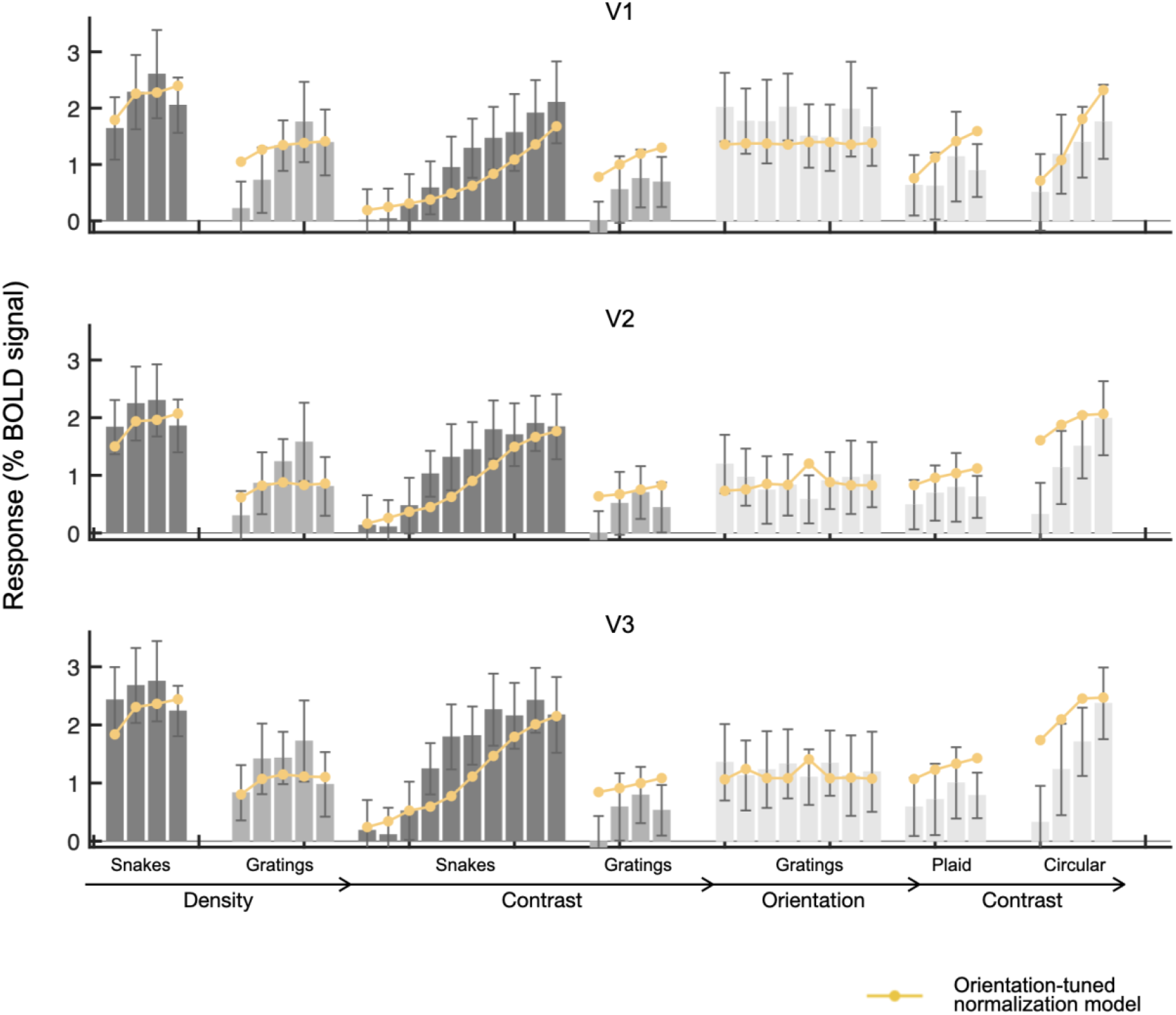
Responses and Figure S6c: Responses and orientation tuned normalization model fits for all stimuli, data set 4

**Figure S6d:**
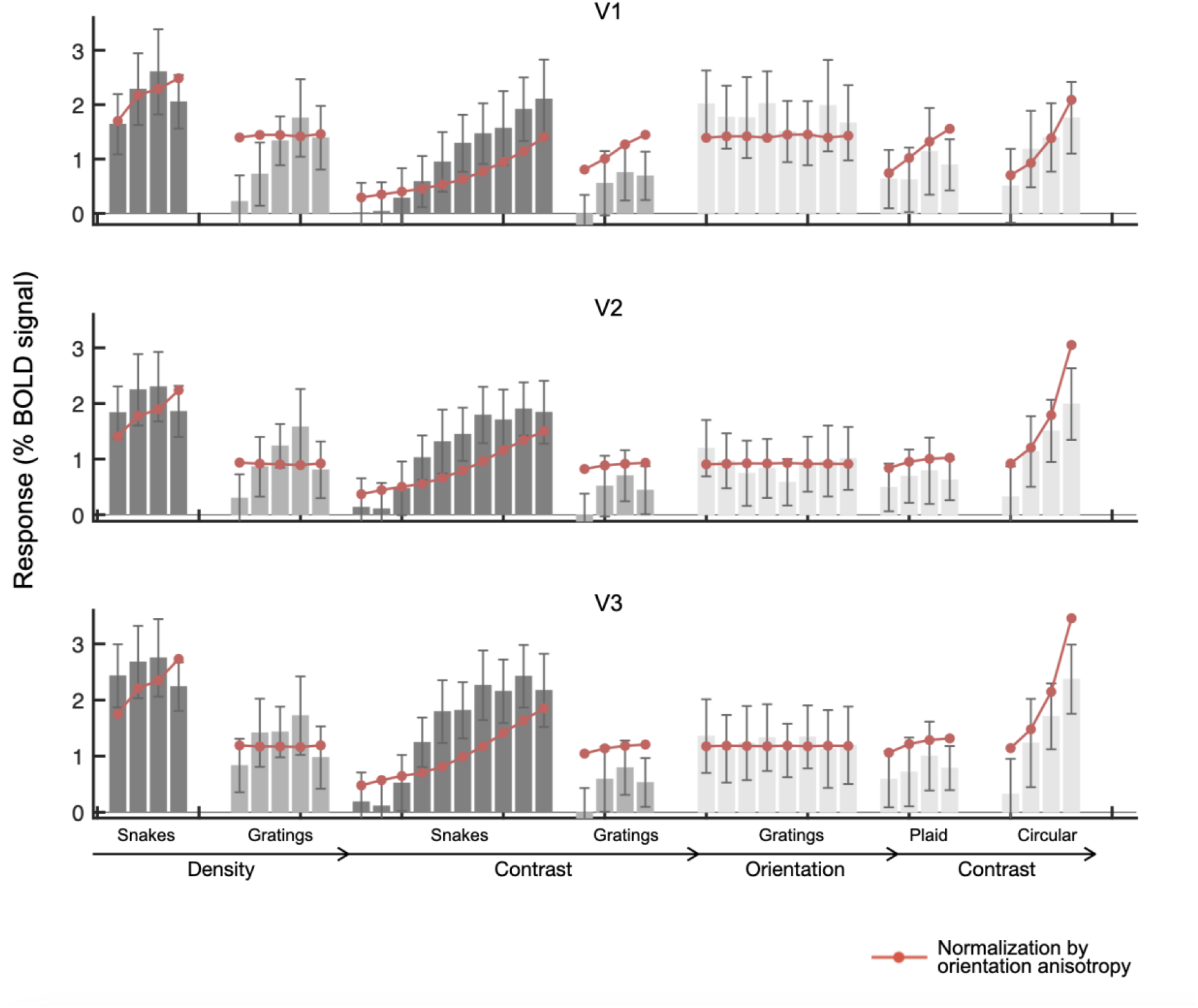
Responses and normalization by orientation anisotropy model fits for all stimuli, data set 4

**Table S1:**
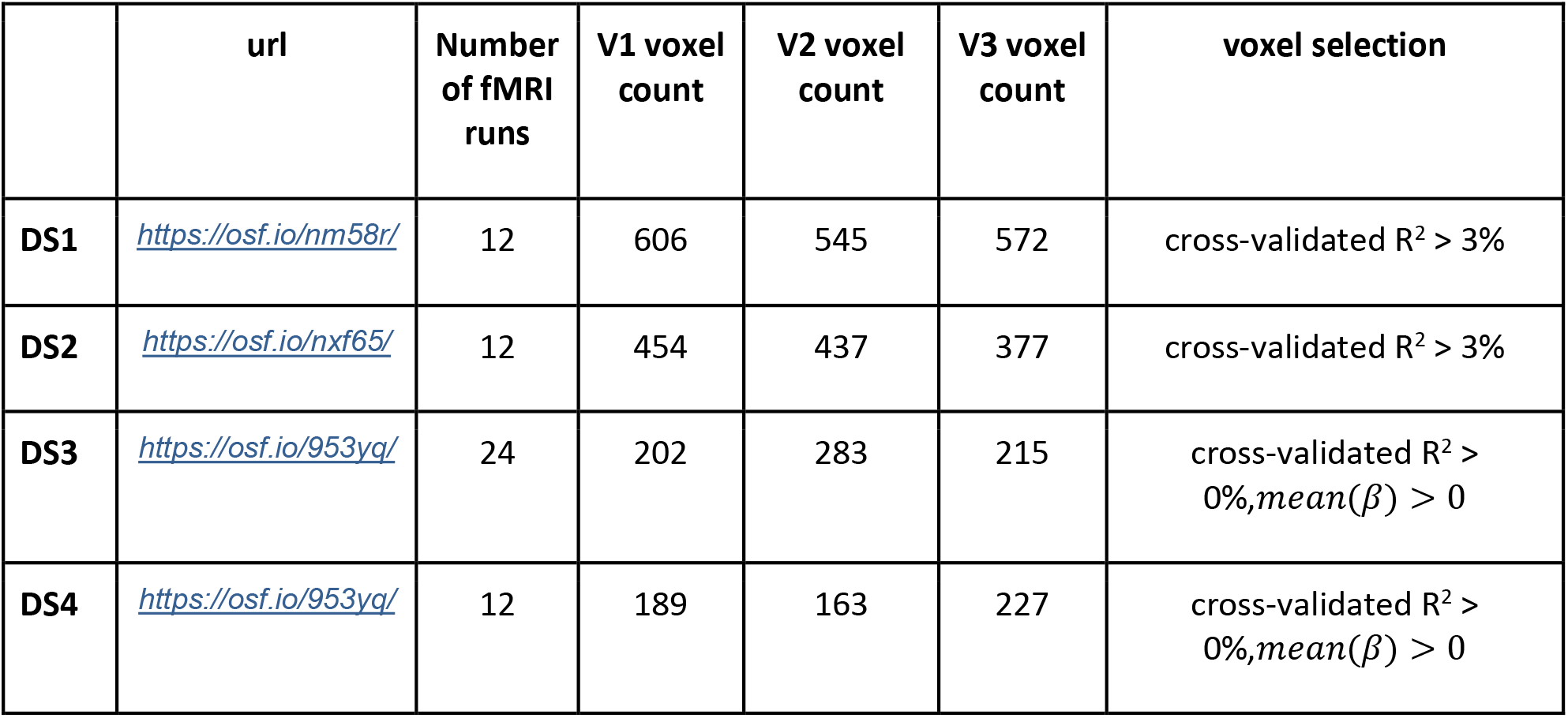
Data set properties for all experiments

**Table S2:** Stimulus set properties for all experiments

|  | url | Size (deg) | Size (pixels, shown to subjects) | Size (pixels, for analysis, excluding padding) | Spatial frequency (cpd) | Number of stimulus classes |
| --- | --- | --- | --- | --- | --- | --- |
| DS1 | https://osf.io/mbtf6/ | $12.5 \times 12.5$ | $400 \times 400$ | $150 \times 150$ | 3 | 50 |
| DS2 | https://osf.io/juya8/ | $18.75 \times 18.75$ | $600 \times 600$ | $225 \times 225$ | 3 | 48 |
| DS3 | https://osf.io/d2r8n/ | $12.5 \times 12.5$ | $800 \times 800$ | $150 \times 150$ | 3 | 39 |
| DS4 | https://osf.io/d2r8n/ | $12.5 \times 12.5$ | $800 \times 800$ | $150 \times 150$ | 3 | 39 |

**Table S3:** Stimulus set group for all experiments

|  | This paper |  | Kay et al 2013 |  |
| --- | --- | --- | --- | --- |
|  | OSF url |  | http://kendrickkay.net/socmodel/stimuli.mat |  |
|  | Number | Description | Number | Description |
| DS1 | 1 2 3 4 5 | SNAKES (density) | Not part of Kay et al. |  |
|  | 47 48 3 49 50 | SNAKES (contrast) |  |  |
|  | 6 7 8 9 10 | GRATINGS (density) |  |  |
|  | 35 36 8 37 38 | GRATINGS (contrast) |  |  |
|  | 11 12 13 14 15 | NOISE BARS (density) |  |  |
|  | 16 17 18 19 20 21 | WAVES (density) |  |  |
|  | 8 22 23 24 | GRATINGS (orientation) |  |  |
|  | 13 25 26 27 | NOISE BARS (orientation) |  |  |
|  | 20 28 29 30 | WAVES (orientation) |  |  |
|  | 31 32 33 34 | GRATINGS (spacing) |  |  |
|  | 39 40 13 41 42 | NOISE BARS (contrast) |  |  |
|  | 43 44 18 45 46 | WAVES (contrast) |  |  |
| DS2 | 1 2 3 4 5 | SNAKES (density) | Not part of Kay et al. |  |
|  | 45 46 2 47 48 | SNAKES (contrast) |  |  |
|  | 6 7 8 9 10 | GRATINGS (density) |  |  |
|  | 33 34 7 35 36 | GRATINGS (contrast) |  |  |
|  | 11 12 13 14 15 | NOISE BARS (density) |  |  |
|  | 16 17 18 19 | WAVES (density) |  |  |
|  | 8 20 21 22 | GRATINGS (orientation) |  |  |
|  | 13 23 24 25 | NOISE BARS (orientation) |  |  |
|  | 18 26 27 28 | WAVES (orientation) |  |  |
|  | 29 30 31 32 | GRATINGS (cross) |  |  |
|  | 37 38 13 39 40 | NOISE BARS (contrast) |  |  |
|  | 41 42 18 43 44 | WAVES (contrast) |  |  |
| DS3,<br>DS4 | 36 37 38 39 | SNAKES (density) | 181-184 | SEPARATION |
|  | 21 22 23 24 25 26<br>27 28 29 30 | SNAKES (contrast) | 159-168 | CONTRAST |
|  | 31 32 33 34 35 | GRATINGS (density) | 176-180 | not used |
|  | 9 10 11 12 1 | GRATINGS (contrast) | 147-150 | GRATING |
|  | 1 2 3 4 5 6 7 8 | GRATINGS (orientation) | 139-146 | ORIENTATION |
|  | 13 14 15 16 | PLAID (contrast) | 151-154 | PLAID |
|  | 17 18 19 20 | CIRCULAR (contrast) | 155-158 | CIRCULAR |
|  | NA | not used | 70-138 | SPACE |
|  | NA | not used | 169-175 | not used (objects/scenes) |
|  | NA | not used | 185-225 | not used (mixtures) |

**Table S4:** **R^2^ Tables**

| *A. Target stimuli* |  |  |  |  |  |  |  |  |  |  |  |  |
| --- | --- | --- | --- | --- | --- | --- | --- | --- | --- | --- | --- | --- |
| Model (free<br>params) | V1 |  |  |  | V2 |  |  |  | V3 |  |  |  |
|  | DS1 | DS2 | DS3 | DS4 | DS1 | DS2 | DS3 | DS4 | DS1 | DS2 | DS3 | DS4 |
| CE (2) | -0.110 | 0.385 | -0.178 | -0.083 | -0.161 | 0.087 | -0.236 | -0.245 | -0.176 | -0.115 | -0.234 | -0.260 |
| DN (3) | -0.185 | 0.347 | -0.194 | 0.009 | -0.161 | 0.097 | -0.391 | -0.273 | -0.190 | -0.029 | -0.381 | -0.350 |
| OTN (3) | **0.288** | **0.802** | **0.673** | **0.591** | 0.714 | **0.799** | **0.773** | **0.692** | **0.870** | 0.587 | **0.722** | **0.756** |
| NOA (3) | 0.008 | 0.765 | 0.529 | 0.486 | **0.777** | 0.689 | 0.615 | 0.539 | 0.798 | **0.786** | 0.542 | 0.530 |

| *B. All stimuli* |  |  |  |  |  |  |  |  |  |  |  |  |
| --- | --- | --- | --- | --- | --- | --- | --- | --- | --- | --- | --- | --- |
| Model (free<br>params) | V1 |  |  |  | V2 |  |  |  | V3 |  |  |  |
|  | DS1 | DS2 | DS3 | DS4 | DS1 | DS2 | DS3 | DS4 | DS1 | DS2 | DS3 | DS4 |
| CE (2) | 0.095 | 0.455 | 0.308 | 0.374 | 0.027 | 0.300 | 0.054 | -0.035 | 0.013 | 0.181 | 0.055 | -0.013 |
| DN (3) | 0.233 | 0.492 | 0.358 | 0.369 | 0.156 | 0.410 | 0.265 | 0.039 | 0.107 | 0.326 | 0.199 | 0.038 |
| OTN (3) | **0.548** | **0.754** | **0.564** | **0.566** | **0.622** | **0.675** | **0.610** | **0.510** | **0.609** | **0.602** | **0.561** | **0.513** |
| NOA (3) | 0.432 | 0.643 | 0.507 | 0.490 | 0.445 | 0.546 | 0.492 | 0.463 | 0.516 | 0.478 | 0.405 | 0.445 |

**Table S5:** Fitted parameters

| *A. Target stimuli* |  |  |  |  |  |  |  |  |  |  |  |  |  |
| --- | --- | --- | --- | --- | --- | --- | --- | --- | --- | --- | --- | --- | --- |
|  |  | **V1** |  |  |  | **V2** |  |  |  | **V3** |  |  |  |
|  |  | **DS1** | **DS2** | **DS3** | **DS4** | **DS1** | **DS2** | **DS3** | **DS4** | **DS1** | **DS2** | **DS3** | **DS4** |
| CE | g | 0.742 | 2.380 | 1.270 | 1.800 | 0.813 | 1.150 | 1.390 | 1.500 | 0.855 | 1.050 | 1.420 | 1.820 |
| | $\alpha$ | 0.133 | 0.129 | 0.053 | 0.097 | 0.064 | 0.082 | 0.017 | 0.055 | 0.041 | 0.055 | 0.019 | 0.044 |
| DN | $\sigma$ | 118.00 | 0.126 | 0.023 | 0.013 | 0.032 | 0.034 | 0.010 | 0.009 | 0.028 | 0.007 | 0.013 | 0.012 |
|  | g | 1.010 | 2.320 | 9.080 | 12.800 | 0.870 | 1.160 | 8.590 | 11.000 | 0.901 | 1.170 | 4.450 | 8.420 |
| | $\alpha$ | 0.306 | 0.189 | 0.949 | 1.000 | 0.137 | 0.158 | 0.850 | 0.963 | 0.094 | 0.179 | 0.538 | 0.718 |
| OTN | $\sigma$ | 0.002 | 0.037 | 0.028 | 0.009 | 0.001 | 0.007 | 0.007 | 0.004 | 0.000 | 0.003 | 0.005 | 0.003 |
|  | g | 7.960 | 4.030 | 6.350 | 11.000 | 8.410 | 2.320 | 5.130 | 11.100 | 8.160 | 2.700 | 4.790 | 10.400 |
| | $\alpha$ | 0.989 | 0.341 | 0.566 | 0.709 | 0.856 | 0.367 | 0.453 | 0.750 | 0.803 | 0.428 | 0.428 | 0.647 |
| NOA | $\sigma$ | 0.001 | 0.005 | 0.001 | 0.000 | 0.000 | 0.001 | 0.000 | 0.000 | 0.000 | 0.000 | 0.000 | 0.000 |
|  | g | 0.339 | 1.530 | 0.970 | 1.150 | 0.495 | 0.821 | 1.180 | 1.040 | 0.603 | 0.783 | 1.200 | 1.330 |
| | $\alpha$ | 0.677 | 0.252 | 0.278 | 0.285 | 0.554 | 0.236 | 0.209 | 0.291 | 0.469 | 0.266 | 0.192 | 0.273 |

| *B. All stimuli* |  |  |  |  |  |  |  |  |  |  |  |  |  |
| --- | --- | --- | --- | --- | --- | --- | --- | --- | --- | --- | --- | --- | --- |
|  |  | **V1** |  |  |  | **V2** |  |  |  | **V3** |  |  |  |
|  |  | **DS1** | **DS2** | **DS3** | **DS4** | **DS1** | **DS2** | **DS3** | **DS4** | **DS1** | **DS2** | **DS3** | **DS4** |
| CE | g | 1.490 | 3.430 | 2.120 | 3.140 | 1.300 | 1.560 | 1.790 | 2.320 | 1.270 | 1.330 | 1.750 | 2.580 |
| | $\alpha$ | 0.406 | 0.289 | 0.221 | 0.285 | 0.230 | 0.202 | 0.100 | 0.197 | 0.174 | 0.144 | 0.088 | 0.158 |
| UN | $\sigma$ | 0.111 | 0.123 | 0.235 | 0.353 | 0.944 | 0.944 | 1.000 | 1.000 | 0.944 | 0.985 | 1.000 | 1.000 |
|  | g | 1.280 | 3.170 | 2.000 | 3.140 | 1.470 | 1.650 | 2.060 | 2.750 | 1.420 | 1.620 | 2.140 | 3.350 |
| | $\alpha$ | 0.216 | 0.159 | 0.116 | 0.157 | 0.144 | 0.118 | 0.072 | 0.125 | 0.111 | 0.107 | 0.076 | 0.120 |
| OTN | $\sigma$ | 357.0 | 82.90 | 403.0 | 403.0 | 403.0 | 348.0 | 403.0 | 403.0 | 403.0 | 168.0 | 399.0 | 403.0 |
|  | g | 233.0 | 9.680 | 67.00 | 117.0 | 34.80 | 10.50 | 10.30 | 50.40 | 18.40 | 3.290 | 6.820 | 30.20 |
| | $\alpha$ | 1.200 | 0.456 | 0.786 | 0.853 | 0.730 | 0.485 | 0.389 | 0.699 | 0.586 | 0.287 | 0.308 | 0.556 |
| NOA | $\sigma$ | 152.0 | 63.60 | 222.0 | 406.0 | 681.0 | 135.0 | 542.0 | 782.0 | 678.0 | 340.0 | 766.0 | 859.0 |
|  | g | 23.20 | 6.390 | 12.70 | 23.30 | 77.90 | 4.130 | 11.80 | 36.00 | 40.40 | 6.820 | 11.60 | 38.60 |
| | $\alpha$ | 0.491 | 0.214 | 0.280 | 0.291 | 0.451 | 0.200 | 0.212 | 0.306 | 0.373 | 0.220 | 0.197 | 0.287 |

